# Comparative characterization of Cas12f orthologs reveals mechanistic features underlying enhanced genome editing efficiency

**DOI:** 10.1101/2025.08.14.670346

**Authors:** Kaoling Guan, Rodrigo Fregoso Ocampo, Paula B. Matheus Carnevali, Cindy J Castelle, Liliana Gonzalez-Osorio, Dominic T. Castanzo, Molly Brothers, Tyler L. Dangerfield, Matthew M. Hooper, Nathan Appleby, Isabella Krudop, Rebecca C. Lamothe, Daniela S. Aliaga Goltsman, Lisa M. Alexander, Cristina N. Butterfield, Kenneth A. Johnson, Christopher T. Brown, David W. Taylor

## Abstract

Miniature CRISPR-Cas12f nucleases are attractive candidates for therapeutic genome editing owing to their compact size and compatibility with adeno-associated virus (AAV) delivery. However, editing efficiencies in mammalian cells are lower than those of larger systems such as Cas12a and SpCas9. The extensive phylogenetic diversity of Cas12f suggests unexplored mechanistic variation with the potential for optimization. Here, we characterize a naturally occurring Cas12f ortholog discovered through metagenomics, Cas12f-MG119-28, which supports robust genome editing in human cells. Through structural, biochemical, and kinetic analyses, we compare Cas12f-MG119-28 with two recently described orthologs, Oscillibacter sp. Cas12f (OsCas12f) and Ruminiclostridium herbifermentans Cas12f (RhCas12f). These orthologs present divergent architectures and regulatory features governing PAM recognition, gRNA binding, dimerization, and DNA cleavage. Notably, Cas12f-MG119-28 achieves efficient R-loop formation via a stable dimer interface and a naturally optimized guide RNA. These discoveries elucidate key mechanistic determinants of Cas12f activity and may offer a framework for engineering compact genome editors with therapeutic potential.

## Introduction

CRISPR-Cas (Clustered Regularly Interspaced Short Palindromic Repeats and CRISPR-associated proteins) systems that provide adaptive immunity against mobile genetic elements in prokaryotes have been repurposed as genome editing tools in various organisms ^1–5^. These highly diverse CRISPR-Cas systems are divided into two main classes (class 1 and class 2), each containing several distinct types ^6^. Class 2 effectors, such as Cas9 (Type II), Cas12 (Type V), and Cas13 (Type VI), use a single effector protein as a nuclease. Among these, *Streptococcus pyogenes* (SpCas9) and *Acidaminococcus sp.* (AsCas12a) have robust nuclease activity and are widely used for genome engineering ^1,7–10^. However, both proteins are relatively large (exceeding 1,300 amino acids), which presents challenges for packaging into adeno-associated viruses (AAV) vectors and mRNA manufacturing, resulting in inefficient delivery and limiting their therapeutic applications ^11^.

To overcome limitations in ribonucleoprotein (RNP) packaging and delivery, smaller CRISPR-Cas subtypes have emerged as candidates for genome editing ^12–14^. One such subtype, type V-F (Cas12f) nucleases, (ranging from 400 – 700 amino acids) ^15^, target double-stranded DNA (dsDNA) with 5’ T-rich or C-rich protospacer adjacent motif (PAM) preferences ^13,16,17^. Cryo- electron microscopy (cryo-EM) studies revealed that Cas12f functions as an asymmetric homodimer, where one monomer cleaves both the target and non-target DNA strands, while the other remains inactive ^18–23^. Although these proteins have yet to reach saturating levels of editing in mammalian cells, rational engineering of Cas12f proteins, as well as their guide RNA (gRNA) scaffolds, have significantly enhanced their gene-editing activity ^21,23–25^.

Continuous discovery of new Cas12f variants underscores the diversity of type V-F nucleases and the potential for uncovering additional naturally optimized orthologs with high intrinsic activity. In this study, we present the structure of a newly identified Cas12f ortholog, Cas12f-MG119-28, which exhibits high editing efficiency in human cells. To contextualize the properties of Cas12f-MG119-28, we solved the structures of OsCas12f and RhCas12f, conducted kinetic analyses of all three orthologs, and performed direct comparisons of structural and biochemical features. We found distinct gRNA binding, PAM recognition and DNA targeting mechanisms. Comparison with other Cas12f nucleases establishes new insights into the structural and mechanistic diversity of these type V-F CRISPR-Cas enzymes.

## Results

### Cas12f-MG119-28 is a highly active CRISPR-Cas nuclease

To explore the diversity of type V nucleases we mined a large database of high-quality assembled microbial metagenomes from diverse environments, uncovering thousands of genes encoding small nucleases in the vicinity of CRISPR arrays. Although divergent, some of these nucleases are related to other Cas12f nucleases when placed in a phylogenetic tree of archetypal type V nucleases (**Fig. 1a**). Newly identified representative sequences Cas12f-MG119-1, Cas12f- MG119-2, Cas12f-MG119-3 and Cas12f-MG119-28 share 28-52% average amino acid sequence identity with reference sequences and range between 433 and 488 amino acids in length (**Fig. 1a, Suppl. Data 1**). Upon inspection of the genomic regions encoding the nuclease genes, we identified the corresponding tracrRNAs and crRNAs. Cas12f-MG119 representatives require large sgRNAs (129 - 156 nt) as empirically confirmed (**Fig. 1b, Suppl. Fig. 1 and Suppl. Data 2**). We determined that these systems recognize T-rich PAMs using an in vitro cleavage assay of an 8N PAM library (**Fig. 1c**), and cut the target strand at 20, 22, and 23 nt from the 5’ PAM sequence and the non-target strand at 11 and 13 nt (**Suppl. Fig. 2** **and 3**). Size exclusion chromatography (SEC) of the purified protein revealed that Cas12f-MG119-28 exists as an obligate dimer even in the absence of any guide RNA (**Suppl. Fig. 4**).

**Figure 1.**
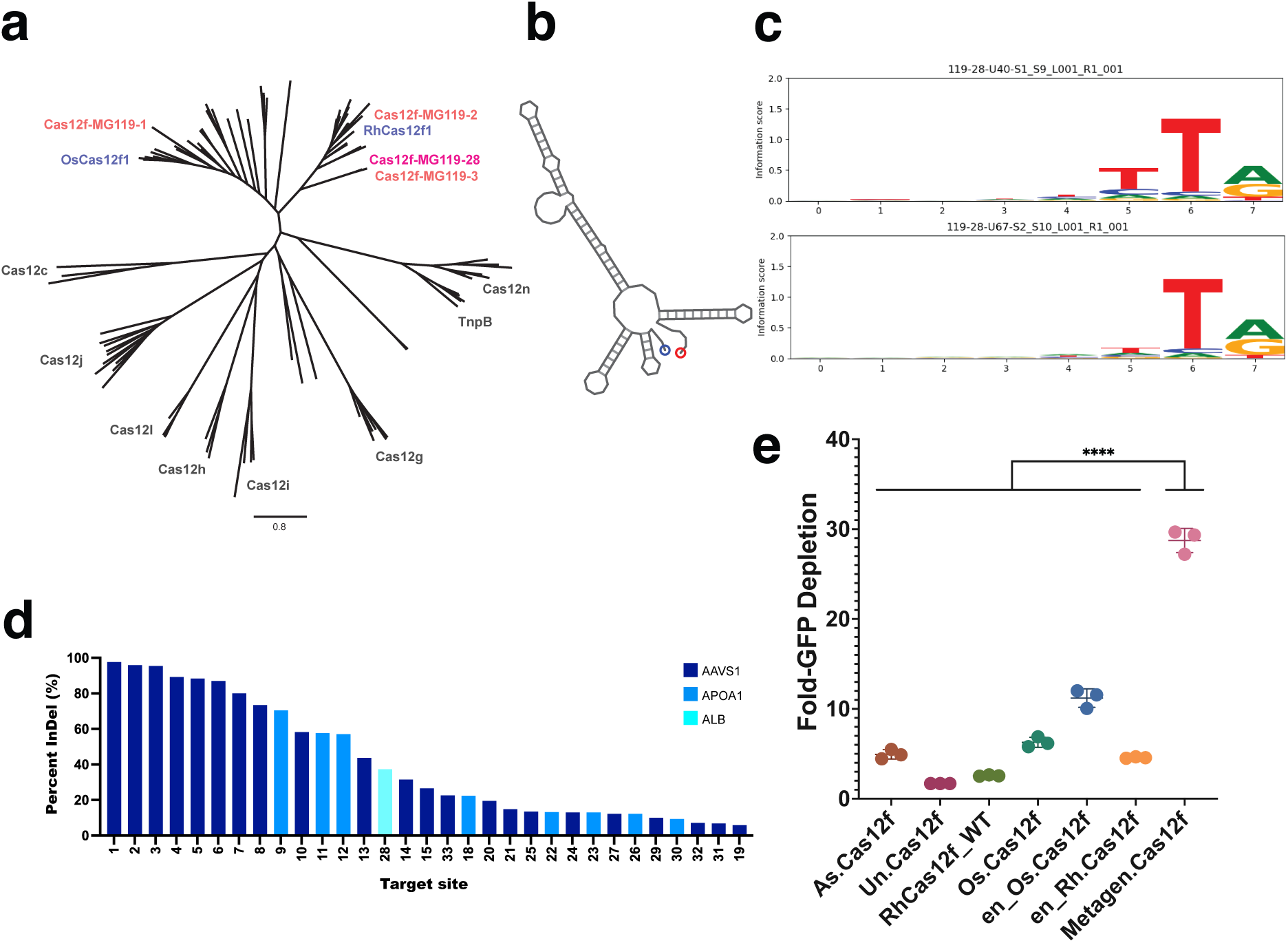
Discovery and characterization of the compact type V Cas12f-MG119-28 nuclease system. **a.** Phylogenetic tree showing representative members of the Cas12f-MG119 family of compact type V systems in relation to other Cas12 and TnpB nucleases. **b.** gRNA (134 nt) secondary structure (Vienna RNAFold). **c.** Sequence logos representing the PAM recognized by Cas12f- MG119-28 across two different spacers. **d.** Guide screen of Cas12f-MG119-28 across multiple loci in K562 cells. Editing efficiency represented as percent of insertion and deletions (InDel) detected in NGS reads. Shown are guides displaying 5% InDels or more for guides targeting the human AAVS1 locus (dark blue), human APOA1 locus (light blue), and human albumin locus (teal). **e.** Comparison of cleavage activity with Cas12f orthologs indicates that Cas12f-MG119- 28 outperforms wildtype and engineered variants of OsCas12f, AsCas12f, and RhCas12f in GFP depletion assays in *E. coli*.

While studying the activity of Cas12f-MG119-28 in human cells, saturating levels of editing efficiency were observed. Results from a guide RNA screen targeting intron 1 of the human *ALB* gene, exon 3 of the human *APOA1* gene, and *AAVS1* showed that 25 target sites across all three loci displayed >10% editing and 12 displayed >50% editing (assessed as the percentage of reads from next-generation amplicon sequencing that contained insertions or deletions). These results, combined with the observation that two target sites achieved >90% editing at *AAVS1* (**Fig. 1d**), suggest that this nuclease is highly active in its native form. To further validate the high levels of activity observed in mammalian cells, we compared the activity of Cas12f-MG119-28 relative to other highly characterized Cas12f orthologs using a GFP reporter assay in *E. coli*. Cas12f-MG119-28 significantly outperformed all orthologs, including engineered variants which have been shown to exhibit high levels of genome editing activity in cells (**Fig. 1e**). Results indicate that this WT nuclease is a valuable candidate for structural and biochemical characterization relative to other Cas12f systems.

### Cryo-EM structures of Cas12f-MG119-28, OsCas12f, and RhCas12f ternary complexes reveal differences in dimer interface

To understand the structural basis for enhanced activity of Cas12f-MG119-28 relative to other orthologs, we first solved a cryo-EM structure of the enzyme bound to its gRNA targeting a 55 bp target DNA. The Cas12f-MG119-28 structures revealed an asymmetric homodimer in a post-cleavage state. The dimer is anchored by multiple interactions within its REC domains as well as direct base stacking interactions with the gRNA scaffold and the R-loop. Consistent with previously solved Cas12f structures, the model revealed a bilobed ternary RNP complex with the REC lobe comprising the recognition (REC) and wedge (WED) domains, while the NUC lobe contains both RuvC and ZF domains. (**Fig. 2a-b, Suppl. Fig. 6, Suppl. Fig. 7**). This enzyme most closely resembles the structure of *Clostridium novyi* (CnCas12f), with REC domains comprised of large extended alpha helices ^22^. However, there are notable differences between the structures of these enzymes: Cas12f-MG119-28 REC domains directly contact stem 2 of its gRNA, as opposed to CnCas12f, which has a shorter stem 2. Our structure contains a fully formed R-loop with both RuvC domains docked, as opposed to the pre-cleavage complex observed in CnCas12f, which forms a partial R-loop. Because of these differences, we sought to obtain a better mechanistic understanding of Cas12f rearrangements as a function of R-loop formation.

**Figure 2.**
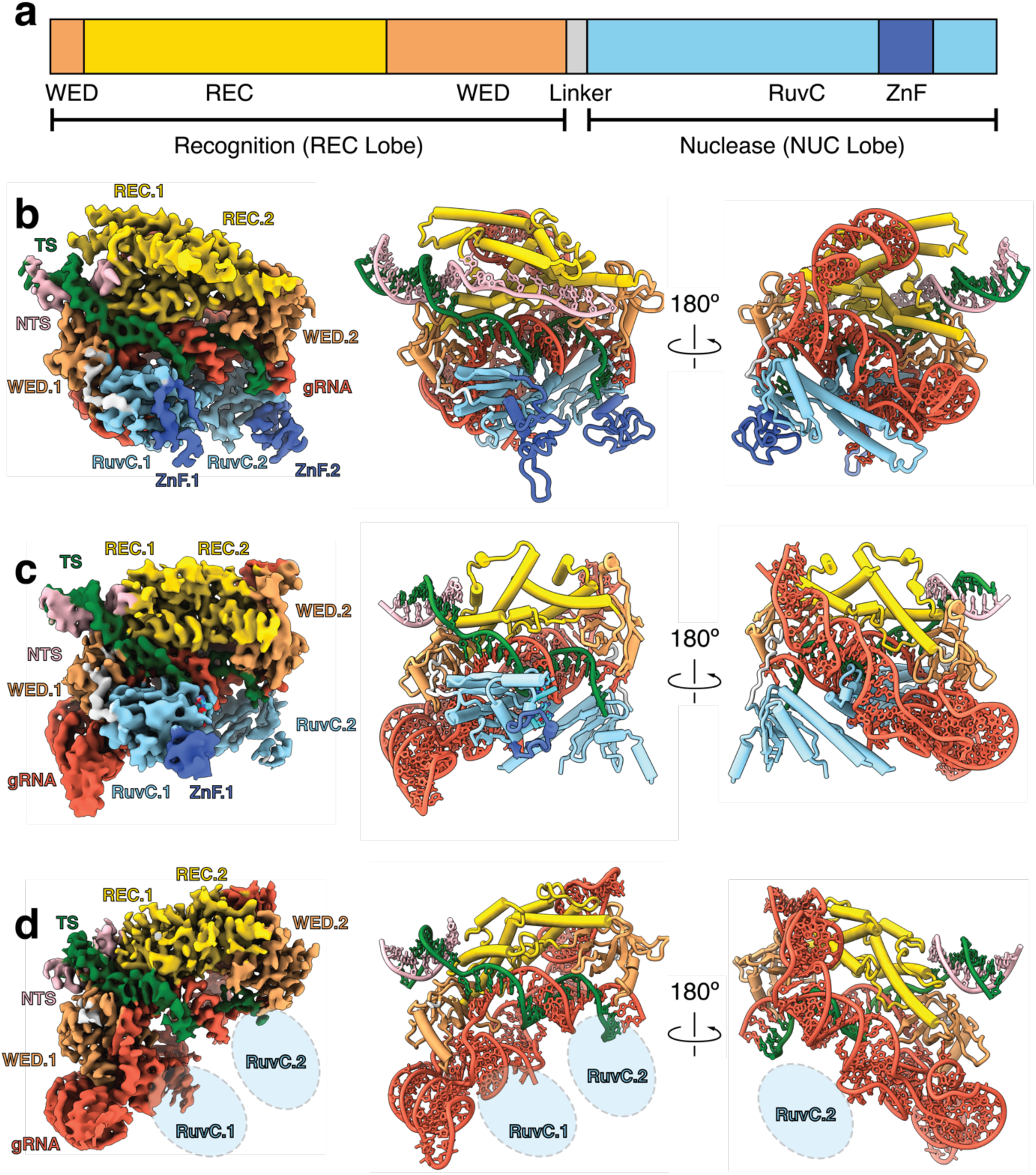
Cryo-EM structure of Cas12f orthologs reveal distinct dimerization interface and RuvC plasticity. **a.** Domain organization of Cas12f. **b-d:** Cryo-EM map (left); front and back views of model (Middle and Right) of Cas12f with a 20-bp R-loop for Cas12f-MG119-28 (**b**); OsCas12f (**c**); and RhCas12f (**d**). Density is colored based on proximity to modeled domains. gRNA is colored red, REC domains are colored gold, WED domains are colored sandy brown, RuvC domains are colored sky blue and ZnF domains are colored royal blue. DNA target and non-target strands are colored forest green and plum, respectively.

Since there are limited Cas12f structures available, we aimed to perform a more thorough comparison of several orthologs known to have high levels of genome editing activity, obtaining structures of OsCas12f and RhCas12f bound to their respective guides and target DNA (**Figure 2c-d** and **Suppl. Fig. 6, Suppl. Fig. 7**). While all structures showed bilobed asymmetric homodimers with a fully formed 20bp R-loop, RhCas12f and OsCas12f contained significantly smaller REC domains relative to Cas12f-MG119-28 (**Fig. 2c-d**). In addition, only Cas12f-MG119- 28 and OsCas12f had resolvable RuvC domains tethered to the R-loop. Surprisingly, neither RuvC domain was resolved in the RhCas12f structure (**Fig. 2d**). These observations suggest that dimerization interfaces vary widely among Cas12f nucleases and that the RuvC domains can sample multiple conformational states and can exhibit different levels of structural plasticity across Cas12f orthologs.

### Cas12f gRNAs exhibit structural and functional diversity

The Cas12f ternary complexes demonstrate significant structural diversity among the cognate gRNA of these nucleases. The gRNA scaffolds of OsCas12f and RhCas12f share a conserved L-shaped architecture (**Fig. 3a-b**), like AsCas12f. However, their RNA is much shorter than their counterpart since they contain only four stem loops, as opposed to the five previously described in AsCas12f (**Suppl. Fig. 8**).

**Figure 3.**
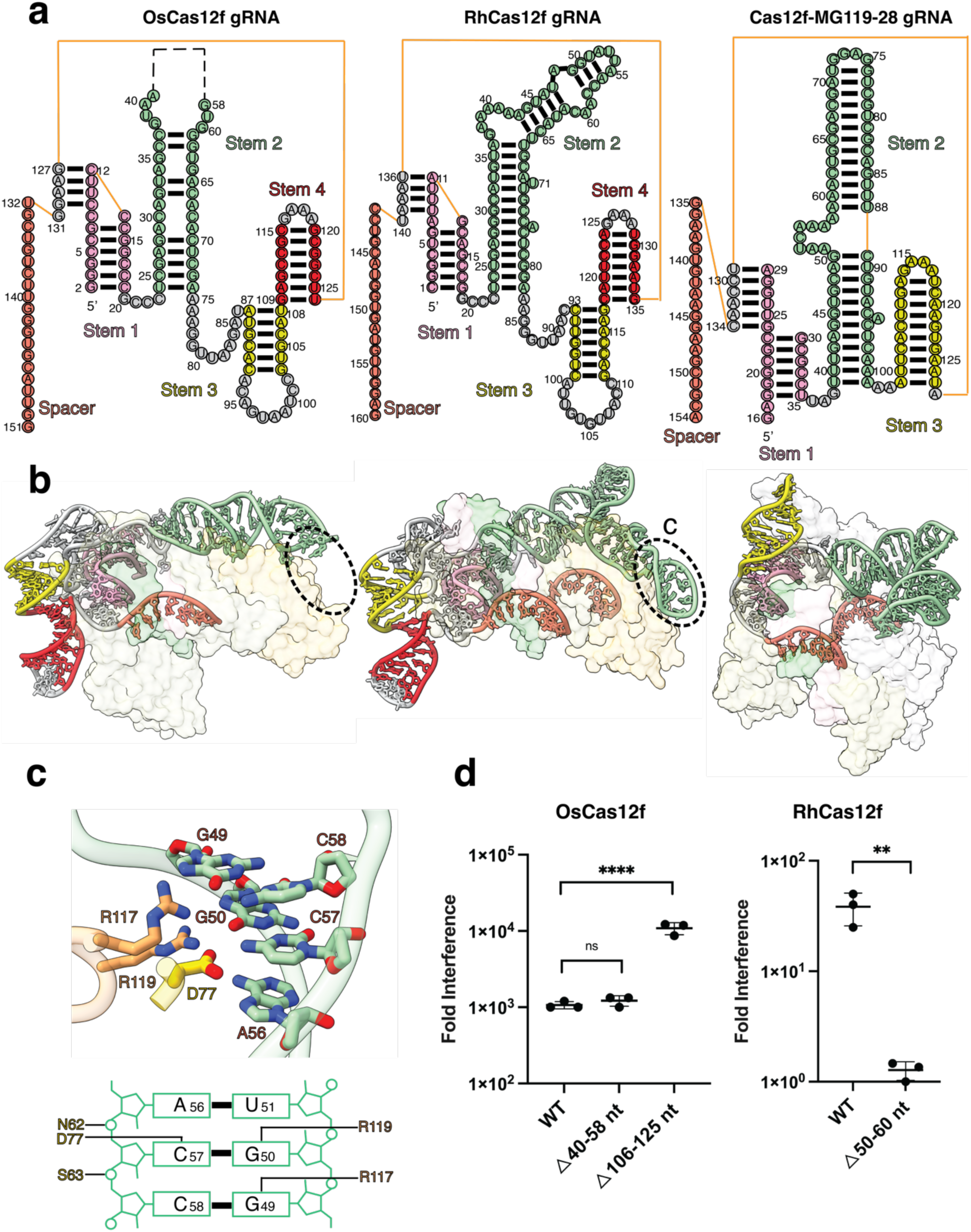
Overview of gRNA architectures in OsCas12f, RhCas12f, and Cas12f-MG119-28 structures. **a.** Schematic diagram of gRNA scaffold of OsCas12f, RhCas12f, and Cas12f-MG119-28. Stems and spacer are labeled and colored accordingly. Unresolved regions are indicated by a dashed line. **b.** Surface views of OsCas12f, RhCas12f, and Cas12f-MG119-28 bound to gRNA. **c.** Model representation of interactions between the RhCas12f WED-REC domain with Stem 2 of its cognate gRNA **d.** Plasmid interference activity of WT and gRNA truncation mutants in OsCas12f and RhCas12f. Data represent mean ± SD (n = 3 biological replicates). Significance was determined by one-way ANOVA or unpaired t-test ns, not significant; **p ≤ 0.01, ****p ≤ 0.0001.

The long Stem 2 of the gRNA of RhCas12f and OsCas12f is nearly perpendicular relative to Stems 3 and 4. While this region is not well resolved for OsCas2f, Stem 2 in the RhCas12f gRNA exhibits a 90° kink that creates additional stabilizing contacts with the Cas12f.2 monomer, making it highly resolvable in our structure (**Fig. 3b**). This region docks into the groove formed by the REC and WED domains and contacts the PAM-interacting residues of monomer 2. Notably this region of the gRNA shares the same sequence with its PAM and is recognized by the same residues involved in PAM binding monomer 1. Specifically, R117 and R119 interact with G49 and G50, respectively, while D77 forms a hydrogen bond with the C57 (**Fig. 3c, Suppl Fig. 9b**). These observations reveal that the same residues in each RhCas12f monomer facilitate dual recognition of the PAM and gRNA, respectively, and prevent non-productive PAM binding by the wrong monomer.

To examine whether the distal end of Stem 2 is necessary for Cas12f activity, we deleted this region in the gRNA of OsCas12f (△40-58 nt) and RhCas12f (△50-60 nt) and assessed their activity in a plasmid interference assay. Deletion of the flexible region in Stem 2 did not affect OsCas12f activity. In contrast, RhCas12f activity was significantly reduced (**Fig. 3d**), indicating that the interaction between Cas12f.2 and Stem 2 of the gRNA is essential for RhCas12f activity. Therefore, engineering of stem 2 in Cas12f gRNAs could provide a strategy towards creating scaffolds to prevent the formation of non-productive conformations of these enzymes.

The Cas12f-MG119-28 gRNA adopts a more compact structure relative to OsCas12f and RhCas12f. Cas12f-MG119-28 gRNA does not contain a Stem 4 region, while its Stem 2 adopts a kinked conformation which allows Stem 2 to dock into the REC lobe of the protein (**Fig. 3b**). Since our structures indicated that the Stem 4 in OsCas12f and RhCas12f did not form direct contacts with the protein, we speculated that deletion of this region could enhance the efficiency of these enzymes. Indeed, deletion of the Stem 4 region (△106-125 nt) in OsCas12f showed a 10-fold increase in plasmid interference (**Fig. 3d**). Cas12f-MG119-28 contains a naturally optimized gRNA scaffold and supports previous findings that removal of Stem 4 in Cas12f nucleases similarly enhances nuclease activity ^23^. Overall, our results provide new mechanistic and structural insights into the gRNA configuration and optimization of Cas12f nucleases.

### Variability in the Cas12f dimerization interface explains enhanced activity by Cas12f- MG119-28

Cas12f dimerizes through interactions between the REC domains of its two monomers. OsCas12f and RhCas12f primarily form dimers via two-layer interactions: the top layer involves the first α-helix of the REC domain (α1), while the bottom layer is formed by two loops located below the α1. In contrast, the dimerization interface of Cas12f-MG119-28 is more complex compared to OsCas12f and RhCas12f (**Fig. 4a-d**). The REC domain of Cas12f-MG119-28 is larger, containing an additional 50 – 60 residues that form two α-helices (α2 and α3). These extended helices in REC.1 and REC.2 interact with the NTS DNA and the hairpin in Stem 2 of the gRNA, respectively (**Suppl. Fig. 10a-c**). These REC domain interactions with non-target DNA and gRNA have not been observed for other Cas12f nucleases and may result in enhanced stability of the complex.

**Figure 4.**
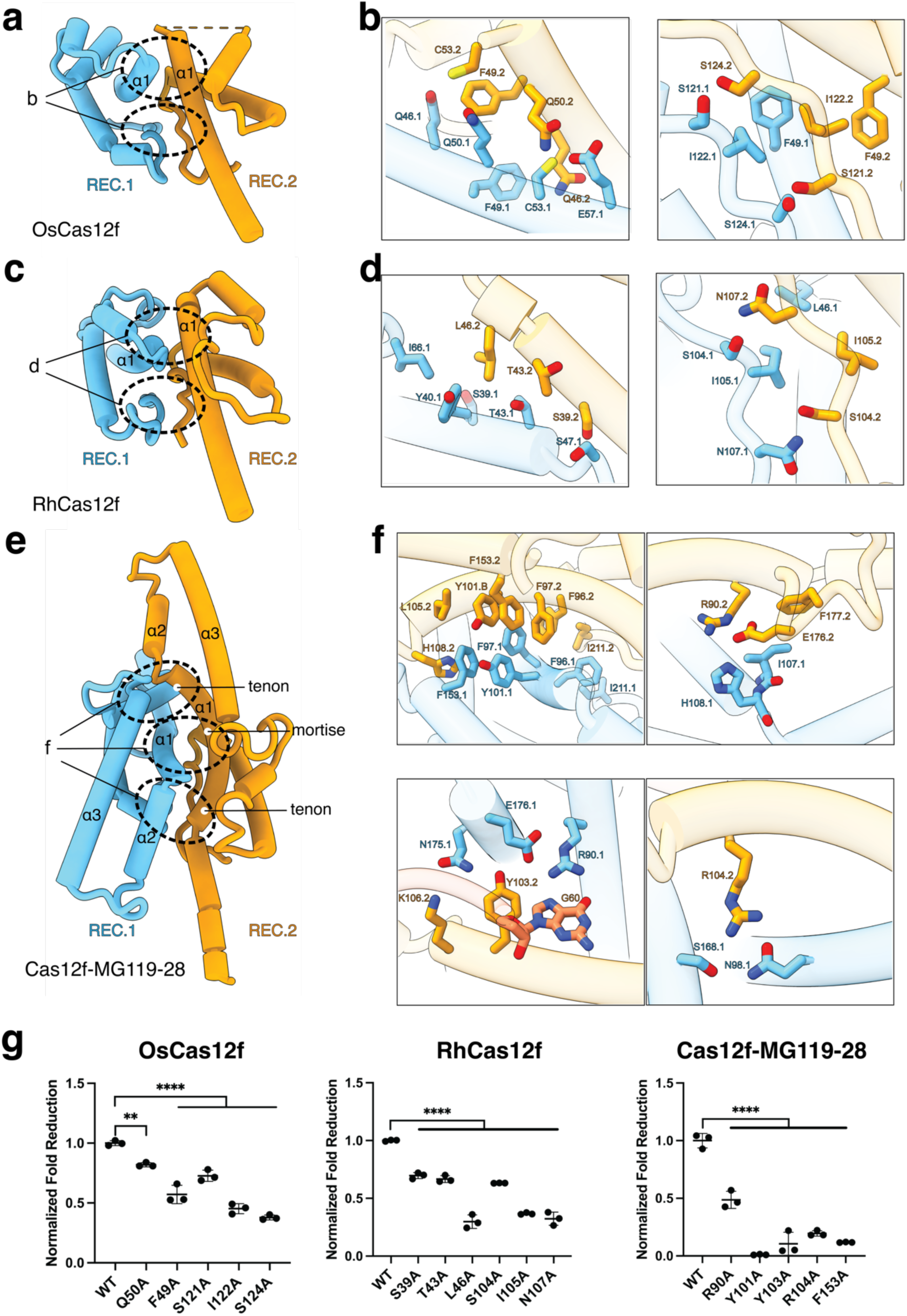
Model representations of REC dimerization interfaces in OsCas12f, RhCas12f, and Cas12f-MG119-28. **a.** Model of OsCas12f dimerization interface **b.** Zoomed in overview of specific interactions that mediate the interactions among the REC interface in OsCas12f. **c.** Model of RhCas12f dimerization interface. **d.** Zoomed in overview of specific interactions that mediate the interactions among the REC interface in RhCas12f **e.** Model of Cas12f-MG119-28 dimerization interface **f.** Zoomed in overview of specific interactions that mediate the interactions among the REC interface in Cas12f-MG119-28. **g.** GFP assays evaluating the effect of different dimerization domain mutants to disrupt specific interactions. Data represent mean ± SD (n = 3 biological replicates). Significance determined by one-way ANOVA: **p ≤ 0.01, ****p ≤ 0.0001.

Cas12f-MG119-28 contains an alpha helix α1 in REC.2 (α1.2) that curves towards REC.1, while the α1 of REC.1 (α1.1) remains mostly linear (**Fig. 4e**). The bending of α1.2 towards α1.1 allows α1.1 to dock onto the α1.2 helix, aided by multiple **π-π** stacking interactions. This unique dimerization mechanism is analogous to a mortise-and-tenon model, where the mortise is the concave slot of the receiving part (α1.2), and the tenon is the protruding structure that fits into the mortise (α1.1) (**Fig. 4f**). Moreover, Cas12f-MG119-28 contains two other mortise-tenon joints located alongside the helices of the REC domain (**Fig. 4f**). The stability at the interface of Cas12f- MG119-28 is mediated through a combination of **π-π** stacking interactions between aromatic residues and Van der Waals interactions among non-polar amino acids. Additionally, salt bridge interactions play a significant role in the stabilization of the dimer (**Fig. 4f**). To validate these observations, we performed alanine substitutions of residues involved in stabilizing the dimerization interface or directly contacting the gRNA or R-loop. These mutations significantly inhibited Cas12f-MG119-28 activity in *E. coli* (**Fig. 4g and Suppl. Fig. 10d**), indicating that dimerization stability mediated by this REC interface is essential for protein function. Overall, our results indicate that a more stable dimerization interface significantly contributes to the enhanced nuclease activity by Cas12f-MG119-28.

### Cas12f intermediate structures reveal differing activation mechanisms for Cas12f variants

To further investigate the molecular mechanisms underlying Cas12f activation, we analyzed our structural datasets for heterogeneity. While the RhCas12f dataset exhibited high structural homogeneity among the selected particles, multiple conformational states were resolved for both OsCas12f and Cas12f-MG119-28.

We solved three distinct structures of OsCas12f in multiple R-loop formation and conformational states (**Fig. 5a-c**). State I showed an 8 bp R-loop with an unresolved RuvC.2, indicating this domain is capable of sampling multiple conformational states, as previously seen for CnCas12f (**Fig. 5a**). State II revealed the RuvC.2 domain docked onto the spacer region of the gRNA prior to R-loop completion, allowing us to model six nucleotides of the gRNA spacer alongside the 8 bp heteroduplex, while RuvC.2 was only partially resolved (**Fig. 5b**). In State III, the structure displayed a fully formed 20 bp R-loop with RuvC.2 domain partially resolved (**Fig. 5c**). Cas12f-MG119-28 displayed two different intermediates, both with a fully formed 20 bp R-loop: a pre-product state prior to RuvC.2 docking (State I) and a post-product state with a fully docked RuvC.2 domain at the PAM-distal end of the spacer (State II) (**Suppl. Fig. 11**).

**Figure 5.**
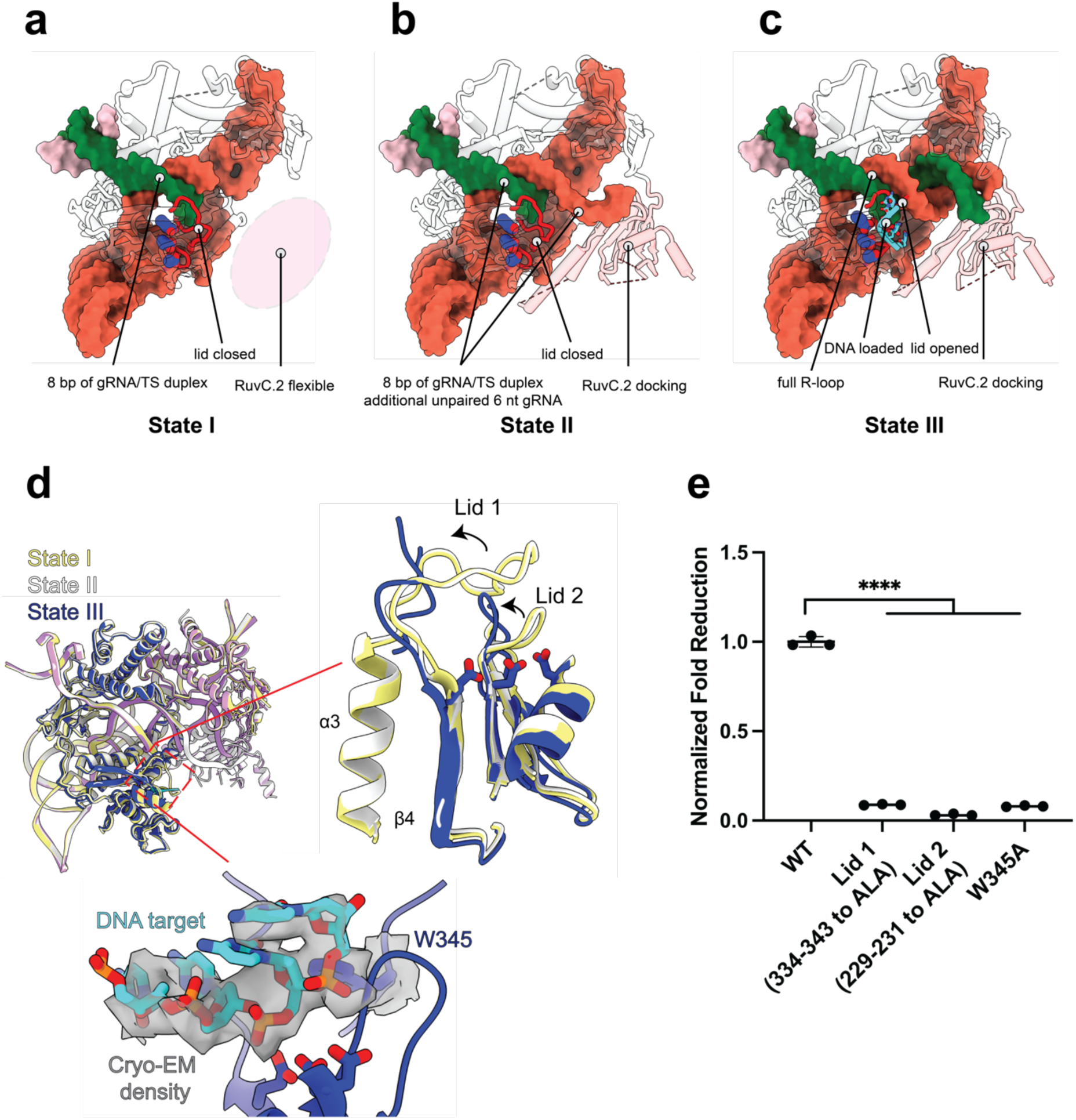
DNA target loading is facilitated by a two-lid opening mechanism. **a-c.** Intermediate state structures illustrating conformational changes at three activation stages of OsCas12f (I-III). Active site residues (blue); lid regions (red). **d.** Structural alignment of intermediate state structures. The region in the dashed square was enlarged to illustrate open-to-close conformational change of two lid regions (black arrow) and the position of the DNA substrate in the catalytic pocket. **e.** GFP depletion assays for lid mutations in OsCas12f.

Although we show that the RuvC.1 domain in OsCas12f is nuclease active and cleaves both strands (**Suppl. Fig. 12**), our structures indicate that RuvC.2 undergoes significant structural rearrangements before eventually docking and stabilizing the PAM-distal region of the R-loop. This enables full R-loop formation in OsCas12f in State II. Conversely, Cas12f-MG119-28 can form a full R-loop prior to RuvC.2 docking. This discrepancy in the order of RuvC docking between OsCas12f and Cas12f-MG119-28 suggests mechanistic differences between these nucleases that may affect their cleavage efficiency.

To understand the importance of these intermediate structures, we examined the RuvC lid in OsCas12f and Cas12f-MG119-28. States I and II in OsCas12f (8bp R-loop formed), show the RuvC lid in a “closed” conformation (**Fig. 5d**). This loop region, connecting the fourth β strand (β4) and the third α-helix (α3) of the RuvC domain, blocks access of the nucleic acid to the catalytic active site, and consequently serves as a crucial regulator of dsDNA cis-cleavage. Upon target loading, this lid has been shown to rearrange to allow for the DNA target strand to pass through to the active site ^18,26–28^. The fully formed R-loop structure in OsCas12f shows that the RuvC.1 lid is in an “open” conformation, indicating that the loop opens only after the R-loop is fully formed. Instead of forming a helix as shown in Un1Cas12f ^18^, the middle region of the lid in OsCas12f State III structure is flexible, and not modeled (D335-L342). We show that this lid is essential for cleavage by mutating E334-R343 to alanine, preventing the lid from undergoing the rearrangements necessary for cleavage, and resulting in significantly reduced OsCas12f activity in GFP depletion assays (**Fig. 5e**). In addition to the RuvC loop, we noticed W345 in OsCas12f positions DNA in the active site of the RuvC domain, like what has been observed in UnCas12f.

A W345A substitution significantly hindered its activity in *E. coli*, indicating this residue is essential for activity in OsCas12f (**Fig. 5d, e**). Moreover, we also observed a smaller loop region (229 - 235 aa, denoted as Lid 2) which undergoes similar conformational changes. Alanine substitutions of the 229 – 231 amino acid region abolished OsCas12f activity, indicating an essential role in nuclease activity.

In contrast to OsCas12f, both structures of Cas12f-MG119-28 showed the lid in an open conformation, indicating that Cas12f-MG119-28 can adapt its cleavage-active conformation prior to RuvC.2 docking. Overall, our structures reveal that RuvC.2 docking is essential for dsDNA cleavage for OsCas12f but not Cas12f-MG119-28. As noted earlier, our results suggest that Cas12f nucleases have distinct cleavage mechanisms, modulated by RuvC rearrangements.

### Kinetic analysis of Cas12f variants

To further investigate the functional differences between these Cas12f variants, we used *in vitro* cleavage assays and stopped-flow fluorescence measurements of R-loop formation for a detailed kinetic analysis. First, we measured cleavage of target and non-target strands by mixing guide-RNP with fluorescently labeled DNA to initiate the reaction. While all enzymes were slower than SpCas9, the cleavage rates varied widely between the Cas12f variants. Cas12f- MG119-28 had the fastest cleavage rate and both target strand and non-target strand cleavage went to completion with data that approximate a single exponential function (**Fig. 6**). Both OsCas12f and RhCas12f displayed highly biphasic behavior with relatively low amplitudes of cleavage, followed by a slow linear approach to completion. This was the most pronounced for RhCas12f and prevented simple interpretation of the results from equation-based fitting.

**Figure 6.**
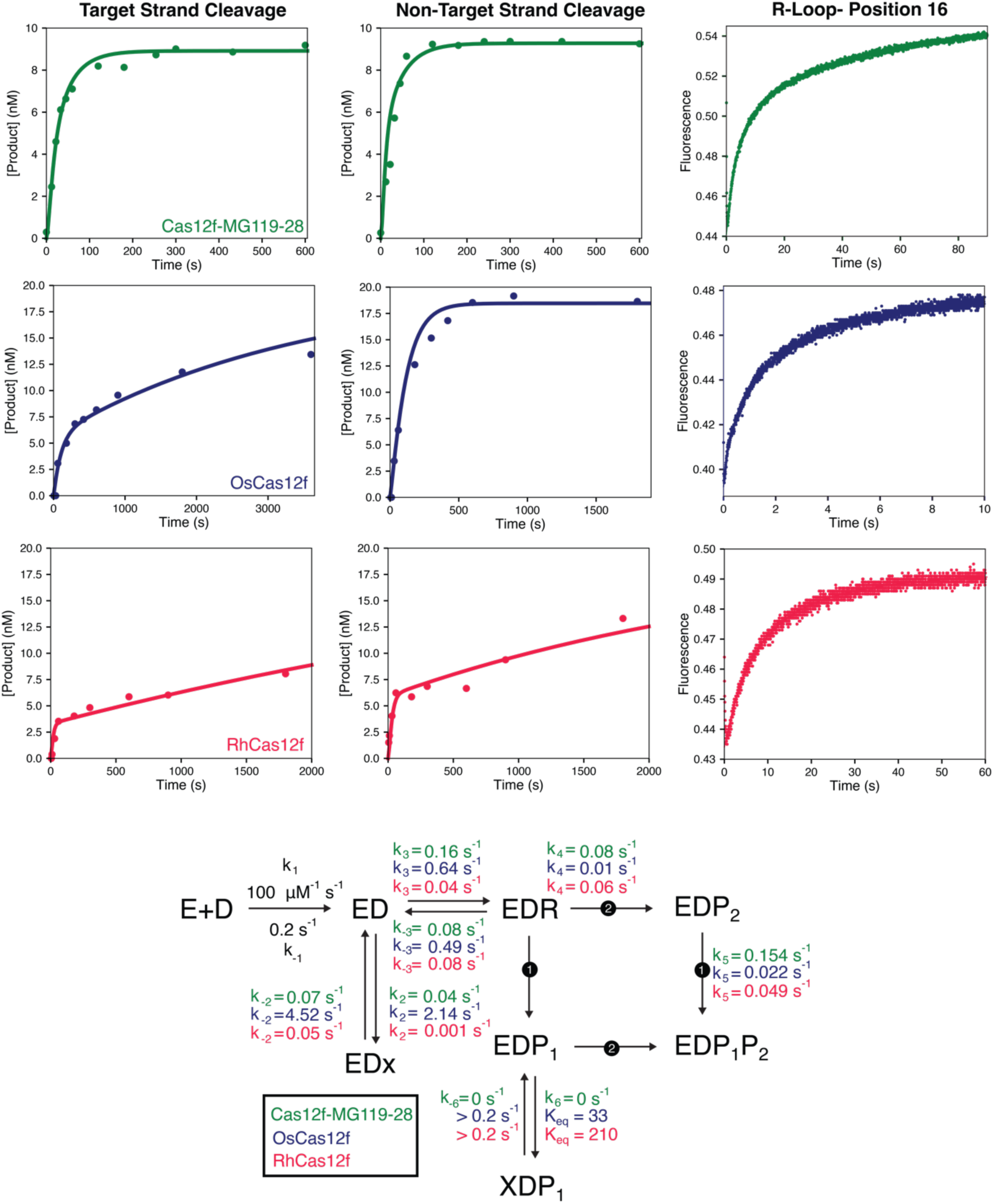
Kinetics of cleavage and R-loop formation by Cas12f variants. **Top:** Cleavage and R-loop formation data for Cas12f-MG119-28, OsCas12f, and RhCas12f are shown in green, blue, and red, respectively. **Bottom:** Kinetic scheme generated in KinTek for DNA binding, R-loop formation, and cleavage for each Cas12f variant. Rate constants in black were locked at the values shown in the fitting.

Next, we measured R-loop formation using a stopped-flow fluorescence assay that we had previously developed for SpCas9 ^29,30^. Data for each variant showed biphasic behavior, indicative of a more complex model than simple DNA binding followed by R-loop formation.

To resolve the complexities in both sets of experiments in the context of a unifying model, we fit the data for each variant globally with KinTek Explorer simulation software ^31,32^. Starting concentrations for each experiment and output observables were entered into the software, allowing us to test multiple models for Cas12f catalysis. Our model has a few unique features for these variants relative to previous models we have published for SpCas9 to account for the complexities in the data. A non-productive state before R-loop formation (EDx) has been added that is accessible from the ED state to account for the biphasic R-loop formation rate (**Fig. 6**). We observed a similar state for Cas9d which was required to account for the biphasic R-loop formation data ^33^. The data for the three variants showed a wide range of forward rate constants (0.05 to 4.5 s^-1^) and reverse rate constants (0.001 to 2.14 s^-1^). In particular, RhCas12f exits this state quite slowly, giving rise to the slow second phase of the cleavage kinetics for target strand cleavage. A non-productive state after non-target strand cleavage (XDP1) was included that is accessible from the cleaved non-target strand state (EDP1). This was required to account for the highly biphasic cleavage data for OsCas12f and RhCas12f. While this step was required to fit the data, only an equilibrium constant for this step could be obtained since the rates of formation and decay of this state had to be faster than the subsequent cleavage step to properly fit the data (> 0.2 s^-1^). Our intermediate structures of OsCas12f reveal that this enzyme requires the RuvC to dock prior to R-loop completion. Consequently, this EDx state could be attributed to a partially formed R-loop prior to RuvC docking.

From this analysis, we can also quantify the preference for each enzyme to cleave target vs. non target strand using flux analysis on the partitioning of the R-loop formed state (EDH). For Cas12f-MG119-28 and OsCas12f, 65 and 70% of the fully formed R-loop state partitioned towards cleavage of the non-target strand first, respectively. For RhCas12f, approximately 45% partitioned towards non-target strand cleavage, essentially showing little preference towards which strand is cleaved first. The unique feature of this class of enzymes, namely the singular active nuclease domain, likely explains these new kinetic observations. In previous studies, we have performed magnesium-initiated cleavage experiments where the enzyme, guide RNA, and DNA are preincubated in the absence of magnesium and then magnesium is added to initiate the reaction. For these enzymes, the measured rates of cleavage were slower than the measured rates of R-loop formation, indicating that the cleavage step was only partially rate-limited by R-loop formation and thus, we were able to extract all rate constants without the need for the magnesium-initiated cleavage experiments.

Global data fitting also resolved rates of R-loop formation for each variant. Forward rates of R-loop formation ranged from 0.04 s^-1^ for RhCas12f to 0.64 s^-1^ for OsCas12f and reverse rates ranged from 0.08 s^-1^ for Cas12f-MG119-28 and Rh-Cas12f to 0.49 s^-1^ for OsCas12f. More importantly, the equilibrium constant for R-loop formation ranged from 0.5 for RhCas12f to 2 for Cas12f-MG119-28. For wild type SpCas9, the equilibrium constant for R-loop formation was measured at 2 ^34^, indicating that R-loop formation for these Cas12f variants is either more reversible or similar to SpCas9 (**Fig. 6** and **Suppl. Fig. 13a-c**).

The higher efficiency and fidelity of these variants can be explained by the *in vitro* kinetics. The reversible R-loop step, coupled to the slower cleavage rates, reflects the general principles seen with higher fidelity SpCas9 variants ^34,35^. That is, with slow cleavage and reversible R-loop formation, off-target DNA has a chance to dissociate before cleavage. Cas12f-MG119-28 is the most efficient at cleaving its target, as it does not get stuck in the EDx or XDP1 state like the other two variants, likely due to full R-loop formation prior to RuvC.2 docking based on our structural studies.

## Discussion

Cas12f nucleases are highly desirable due to their size and levels of genome editing activity. Through mining of a large, assembly-driven metagenomics database, we identified a natural ortholog of Cas12f, Cas12f-MG119-28, that exhibits high levels of activity in vitro and in mammalian cells. We show that this enzyme is advantageous due to its recognition of a minimally restrictive PAM, naturally optimized gRNA, and ability to remain a highly stable dimer.

To better understand how Cas12f-MG119-28 achieves higher levels of editing relative to other Cas12f orthologs, we performed a comprehensive structural comparison across various members of this CRISPR subtype. Prior to this study, three structures of Cas12f nucleases had been described. Here we solved cryo-EM structures of Cas12f-MG119-28 and two additional Cas12f nucleases, also reported to generate high levels of indels in mammalian cells. Structures of Cas12f-MG119-28, OsCas12f, and RhCas12f in combination with AsCas12f, UnCas12f, and CnCas12f, showed significant differences along their dimerization interfaces, which play a significant role in stabilizing the enzyme with Cas12f-MG119-28 having the largest number of interacting residues and interface area (**Suppl. Fig 14**). Interestingly, Hino *et al*. demonstrated AsCas12f activity could be significantly improved by dimerization interface stabilization ^23^. They used unbiased deep mutational scanning which revealed specific dimer residues that increase dimer contacts, bringing engineered AsCas12f closer to the native dimer interface observed in the highly active Cas12f-MG119-28. Additionally, the dimerization interface of Cas12f-MG119- 28 features an extensive contact area forming twice the amount of contacts compared to OsCas12f and RhCas12f, mediated mainly by strong protein-protein interactions between its two large REC domains (**Suppl. Fig. 14**). The dimerization interface is further stabilized by contacts with its gRNA and non-target strand DNA, which forms a highly stable paired region with an optimal size to allow the hairpin to interact with the REC domain, a feature unique to Cas12f-MG119-28. gRNA engineering efforts have shown that enhancing duplex stability by promoting more stable base pairing significantly enhances activity in cells. While we show the gRNA of Cas12f-MG119-28 has already been naturally optimized and contacts the REC domain for additional stability, structures of RhCas12f also revealed that the second hairpin of the gRNA binds the PAM-interacting residues of the non-productive monomer further stabilizing the RNP.

Structures of intermediate states determined in this work show novel conformational states of Cas12f nucleases. Previous studies have shown that CRISPR-Cas systems exhibit extensive domain rearrangements sequentially with the formation of an R-loop ^36^. Notably, intermediate structures of OsCas12f revealed a highly stable dimer throughout the entirety of R- loop formation. These structures showed only RuvC.2 exhibits a high degree of flexibility, while the rest of the protein remains relatively static. In contrast to other Cas systems, where REC domains are used for R-loop stabilization ^36^, OsCas12f uses its RuvC domain to stabilize the PAM-distal end of the spacer and facilitate R-loop formation. However, Cas12f-MG119-28 uses extended REC domains to facilitate additional contacts with the R-loop and the non-target strand, enabling full R-loop formation prior to RuvC docking.

These structural observations are supported by our kinetics studies and models. We defined an EDx state, in which the enzyme is bound to its substrate but preceding completion of R-loop formation. Our kinetic analysis reveals OsCas12f spends significantly longer in this state. This characteristic of OsCas12f can be explained by our 8bp R-loop intermediate structure, necessitating RuvC docking prior to adopting a catalytically active conformation. In contrast, both Cas12f-MG119-28 and RhCas12f do not transition favorably into this EDx state. Congruent with these observations, we were only capable of capturing structures of Cas12f- MG119-28 and RhCas12f with a fully formed R-loop. However, it must be mentioned that while Cas12f-MG119-28 did show RuvC rearranges to bind the PAM-distal end of the R-loop, this might not be necessary for RhCas12f since both RuvC domains were not defined in this structure. Our kinetic studies also reveal that R-loop formation is less reversible for Cas12f- MG119-28, pulling the reaction towards the largely irreversible step of DNA cleavage.

Through a combination of structural, kinetic, and biochemical assays, we highlight multiple mechanistic differences among Cas12f orthologs, and expand our understanding of the structural features important for achieving high nuclease activity. We show that a highly stable dimerization interface in conjunction with a naturally optimized gRNA enables Cas12f-MG119-28 to form an efficient R-loop without requiring prior domain rearrangements. This efficient compact nuclease is a promising candidate for a wide range of biotechnological and therapeutic applications that require compact nucleases for efficient delivery.

## Material and methods

### Sample collection and sequencing

Microbiome samples from abandoned animal stool were collected without disturbing the animals and processed for metagenomic sequencing as described in Goltsman et al ^14^. Other microbiome sequencing data publicly available was downloaded from the NCBI SRA. A list of samples is provided in the Supplementary Information (**Suppl. Data 1**). Raw sequencing reads were trimmed using BBMap (Bushnell B., sourceforge.net/projects/bbmap/), assembled using Megahit ^37^ and genes were predicted using Prodigal ^38^.

### Identification of compact type V nucleases

The discovery of compact type V nucleases was based on homology searches performed using the HMMER software. Type V nuclease sequence hits were retained if they met the following criteria: (i) the hmmsearch e-value was ≤ 10-5, (ii) the genes encoding the nuclease were within 1 Kb from a CRISPR array, and (iii) the amino acid sequence length ranged between 350 and 700 aa. MMSeqs2 (https://github.com/soedinglab/MMseqs2) was used to cluster sequences at 100% amino acid identity, with coverage mode 1 and 80% coverage of the target sequence (parameters --cov-mode 1 -c 0.8 --min-seq-id 1.0).

Sequence representatives and known reference sequences (**Suppl. Data 1 and 3**) were chosen to build a multiple sequence alignment using MAFFT (https://mafft.cbrc.jp/alignment/software/) with the Needleman-Wunsch algorithm for global alignment, and FastTree (https://doi.org/10.1371/journal.pone.0009490) was used to build a phylogenetic tree. Examination of individual clades on the phylogenetic tree, including the nuclease gene’s genomic context, led to the identification of novel Cas12f-M119 sequences.

Although the convention is to name CRISPR nucleases based on the organism that encodes them, it is not possible to do so accurately in cases when the species or strains have not yet been characterized. Therefore, the naming scheme used here is as follows: the type V system most closely related to these sequences (i.e., Cas12f) is followed by the suffix MGX-Y, where MG indicates that the proteins are derived from metagenomic fragments, X represents the family identifier and Y indicates the member identifier. For example, Cas12f-MG119-28, a Cas12f enzyme recovered from metagenomics data, is the 28th member of family 119.

### Search for novel tracrRNA sequences and gRNA design

To identify putative tracrRNA sequences, the intergenic sequence and a minimal array adjacent to the Cas12f-MG119-2 nuclease gene were expressed in transcription-translation reaction mixtures using myTXTL®Sigma 70 Master Mix Kit (Arbor Biosciences). The final reaction mixtures contained 5 nM nuclease DNA template, 12 nM intergenic DNA template, 15 nM minimal array DNA template, 0.1 nM pTXTL-P70a-T7rnap and 1X of myTXTL®Sigma 70 Master Mix. The reactions were incubated at 29 °C for 16 hours then stored at 4 °C. Cleavage activity was confirmed by mixing 5 nM of a target plasmid DNA library representing all possible 8N PAMs, a 5-fold dilution of the TXTL expressions, 10 nM Tris-HCl, 10 nM MgCl2 and, 100 mM NaCl at 37 °C for 2 hours. Reactions were stopped and cleaned with HighPrep™ PCR clean up beads (MAGBIO Genomics, Inc.) and eluted in Tris EDTA pH 8.0 buffer. Cleavage was verified by NGS following the same methods used to determine the PAM sequence.

To obtain the sequence of the tracrRNA and the crRNA, RNA was extracted from TXTL lysate following the Quick-RNA™ Miniprep Kit (Zymo Research) and eluted in 30-50 µL of water. 100 ng - 1 ug of total RNA from each sample were prepped for RNA sequencing using the NEBNext Small RNA Library Prep Set for Illumina (New England Biolabs Inc.). Amplicons between 150-300 bp were quantified by Tapestation and Qubit and pooled to a final concentration of 4 nM. A final concentration of 12.5 pM was loaded into a MiSeq V3 kit and sequenced in a Miseq system (Illumina) for 176 total cycles. Sequencing reads were used to identify the tracrRNA sequence by mapping back to the original DNA fragments encoding the Cas12f-MG119-2 system.

To identify additional non-coding regions containing potential tracrRNAs, the sequence of the active Cas12f-MG119-2 tracrRNA was mapped to other DNA fragments encoding homologs (e.g., Cas12f-MG119-1 and Cas12f-MG119-3). The 3’ end of the predicted tracrRNA sequences as well as the 5’ end of the repeat sequences were trimmed, and then connected with a GAAA tetraloop to generate the gRNAs. The predicted gRNAs were tested in vitro using the same methods used to determine the PAM sequence. The verified sequences for Cas12f- MG119-2 and Cas12f-MG119-3 were used to generate covariance models to predict tracrRNAs for other homologous systems. Covariance models are built from a multiple sequence alignment (MSA) of the active and predicted tracrRNA sequences. The secondary structure of the MSA is obtained with RNAalifold (Vienna Package), and the covariance models are built with Infernal packages (http://eddylab.org/infernal/). Other DNA fragments containing candidate nucleases including Cas12f-MG119-28 were searched using the covariance models with the Infernal command ‘cmsearch’. The gRNAs were tested via in vitro cleavage reactions as described below, and activity was confirmed for a short (134 nt without spacer; **Suppl. Data 2**) and a long version of the gRNA (152 nt without spacer; **Suppl. Data 2; Suppl. Fig. 1**).

### In vitro cleavage reactions to enable PAM determination

5 nM of nuclease amplified DNA templates and 25 nM gRNA amplified DNA templates including one of the following spacer sequences: GTCGAGGCTTGCGACGTGGT (U67 spacer) or TGGAGATATCTTGAACCTTG (U40 spacer) were expressed at 37 °C for 3 hours with PURExpress® In Vitro Protein Synthesis Kit (New England Biolabs Inc.). Plasmid library DNA cleavage reactions were carried out by mixing 5 nM of the target library representing all possible 8N PAMs, a 5-fold dilution of PURExpress expressions, 10 mM Tris-HCl pH 7.9, 10 mM MgCl2, 100 µg/ml BSA, and 50 mM NaCl (NEB 2.1 Buffer, NEB Inc.) at 37 °C for 2 hours. Reactions were stopped and cleaned with HighPrep™ PCR clean up beads (MAGBIO Genomics, Inc.) and eluted in Tris EDTA pH 8.0 buffer.

To determine the PAM sequence and the target strand (TS) cleavage site, 3 nM of the cleavage product ends were blunted with 3.33 µM dNTPs, 1X T4 DNA ligase buffer, and 0.167 U/µL of Klenow Fragment (New England Biolabs Inc.) at 25 °C for 15 minutes. To confirm the PAM sequence and determine the non-target (NTS) strand cleavage site, Mung Bean Nuclease and 1X Mung Bean Nuclease Buffer (New England Biolabs Inc.) was used to blunt the cleavage product ends at 30 °C for 15-30 minutes. 1.5 nM of the cleavage products were ligated with 150 nM adapters, 1 X T4 DNA ligase buffer (New England Biolabs Inc.), 20 U/µL T4 DNA ligase (New England Biolabs Inc.) at room temperature for 20 minutes. The ligated products were amplified by PCR with NGS primers and sequenced by NGS to obtain the PAM sequence. The cut sites on the TS and the NTS were determined from read counts at each nucleotide position. Active systems that successfully cleaved the PAM library yielded a band around 188 or 205 bp in an agarose gel when blunted with Klenow Fragment and a 195 bp band when blunted with Mung Bean Nuclease (**Suppl. Fig. 2**).

### Nuclease mRNA production

The Cas12f-MG119-28 nuclease mRNA sequence was codon optimized for human expression (Twist), then synthesized and cloned into a high copy ampicillin plasmid (Twist Biosciences). Synthesized constructs encoding T7 promoter, UTRs, nuclease ORF, and NLS sequences were digested from the Twist backbone with HindII and BamHI (NEB) and ligated into a pUC19 plasmid backbone with T4 DNA ligase and 1X reaction buffer (NEB). The complete nuclease mRNA plasmid consists of an origin of replication, ampicillin resistance cassette, the synthesized construct, and an encoded polyA tail. Nuclease mRNA was synthesized via in vitro transcription (IVT) using the linearized nuclease mRNA plasmid. This plasmid was linearized by incubation at 37 °C for 16 hours with SapI (NEB) enzyme. The linearization reaction consisted of a 50 uL reaction containing 10 ug pDNA, 50 units Sap I, and 1X reaction buffer. Linearized plasmid was purified with Phenol:Chloroform:Isoamyl Alcohol (25:24:1, v/v), precipitated in EtOH, and resuspended in nuclease free water at an adjusted concentration of 500 ng/uL. The IVT reaction to generate nuclease mRNA was performed at 50°C for 1 hr under the following conditions: 1 ug linearized plasmid; 5 mM ATP, CTP, GTP (NEB), and Nl-methyl pseudo-UTP (TriLink); 18750 U/mL Hi-T7 RNA Polymerase (NEB); 4 mM CleanCap AG (TriLink); 2.5 U/mL Inorganic E. coli pyrophosphatase (NEB); 1000 U/mL murine RNase Inhibitor (NEB); and 1X transcription buffer. After 1 hr, IVT was stopped, and plasmid DNA was digested with the addition of 250 U/mL DNaseI (NEB) and incubated for 10 min at 37 °C. Purification of nuclease mRNA was performed using an RNeasy Maxi Kit (Qiagen) using the standard manufacturer protocol. Transcript concentration was determined by UV (NanoDrop) and further analyzed by capillary gel electrophoresis on a Fragment Analyzer (Agilent).

### Editing efficiency in Mammalian Cells

Experiments were performed in K562 cells grown and passaged in Iscove’s Modified Dulbecco’s Medium (Gibco) supplemented with 10% (v/v) fetal bovine serum at 37 °C with 5% CO2. For nucleofection with Cas12f-MG119-28 mRNA, 500 ng mRNA and 200 pmol of gRNA were mixed together and incubated on ice until cells were prepared. The guide sequences targeting human Albumin intron 1, human APOA1 exon 3, and AAVS1 (**Suppl. Data 2**) were encoded in the longer version of the Cas12f-MG119-28 gRNA (152 nt scaffold).

Approximately 1E+5 cells were transfected with mRNA plus gRNA as recommended by the Amaxa™ 4D-Nucleofector™ Protocol in a 4D-Nucleofector™ System (Lonza). Transfected cells were grown for 3 days, harvested, and gDNA was extracted with QuickExtract (Lucigen) per manufacturer’s instructions. Targeted regions for InDels were amplified using Q5 High-Fidelity DNA polymerase (New England Biolabs) with primers and extracted DNA as the templates. PCR products were purified by HighPrep PCR Clean-up System (MAGBIO) per manufacturer’s instructions. PCR primers appropriate for use in NGS-based DNA sequencing were generated, optimized, and used to amplify the individual target sequences for each guide RNA. The amplicons were sequenced on an Illumina MiSeq machine and analyzed with CRISPResso2 (https://github.com/pinellolab/CRISPResso2) to measure InDel frequency.

### Plasmid construction

The codon-optimized coding sequences of cas12f were synthesized and cloned into a pET-29b expression vector with C-terminal 6xHis tag (Twist Biosciences). DNA templates for gRNA production were synthesized and cloned into a pUC19 vector (Twist Bioscience). Site-directed mutagenesis was performed using Q5 High-Fidelity DNA Polymerase and KLD Enzyme Mix (New England Biolabs). Oligonucleotides were synthesized by Integrated DNA Technologies. All plasmids were verified by whole plasmid sequencing (Plasmidsaurus).

### Protein and gRNA production

Cas12f-MG119-28 proteins were expressed in the pMGBΔ vector, which has the following sequence architecture: 6xHis-(GS)2-3C protease motif-nucleoplasmin bipartite NLS- (GGS)1-(GS)1-Cas12f-MG119-28-(GGS)3-SV40 NLS. E. coli (NEBExpress Iq Competent E. coli, NEB C3037I) cultures were grown at 37 °C in 2xYT media (1.6 % tryptone, 1 % yeast extract, 0.5 % NaCl) or TB media (Teknova T0690) with 100 µg / L Carbenicillin. At OD600 ≈ 0.8 – 1.2, cultures were induced with 0.5 mM IPTG (GoldBio I2481) and incubated at 18°C overnight or 24°C for 4-6 hrs, depending on construct. Cultures were then harvested by centrifugation at 6,000 x g for 10 min, and pellets were resuspended in Nickel_A Buffer (50 mM Tris pH 7.5, 750 mM NaCl, 10 mM MgCl2, 20 mM imidazole, 0.5 mM EDTA, 5 % glycerol, 0.5 mM TCEP) + protease inhibitors (Pierce Protease Inhibitor Tablets, EDTA-free, ThermoFisher A32965) and stored a -80 °C.

Cell pellets were thawed and the volume supplemented to 120 mL with Nickel_A buffer + 0.5 % n-Octyl-ß-D-glucoside detergent (P212121, CI-00234). Samples were sonicated in an ice-water bath at 75% amplitude for a total processing time of 3 min using a 15 s on / 45 s off cycle. Lysates were clarified by centrifugation at 30,000 x g for 25 min, and supernatants batch bound to 5 mL Ni-NTA resin (HisPur Ni-NTA Resin, ThermoFisher 88223) for ≥ 20 min.

Samples were loaded onto a gravity column and washed with 30 CV Nickel_A Buffer, then eluted in 4 CV Nickel_B Buffer (Nickel_A Buffer + 250 mM imidazole) before concentrating in a 50 kDa MWCO concentrator (Amicon Ultra-15, MilliporeSigma UFC9050). Samples were taken throughout the purification process and run on an SDS-PAGE protein gel (BioRad #4568126), which was imaged on a BioRad ChemiDoc in the stain-free channel following 5 min UV activation (Suppl. Fig. 3). Protein samples were incubated with HRV-3C protease (GenScript Z03092) for 15 min at room temperature, then loaded onto an S200i 10 / 300 GL column (Cytiva 28-9909-44) and run into SEC buffer (50mM Tris pH 7.5, 250 mM NaCl, 10 mM MgCl2, 0.5 mM EDTA, 5 % glycerol, 0.5 mM TCEP). Peak fractions were pooled and concentrated in a 50 kDa MWCO concentrator.

Os- and RhCas12f proteins were produced and purified as previously described with minor modifications ^17^. Briefly, protein samples were produced overnight in *Escherichia coli* Nico21(DE3) (New England Biolabs) in the presence of 0.5 mM IPTG at 16 °C. Cells were harvested by centrifugation and lysed by sonication in Lysis buffer (50 mM Tris-HCl pH 8.0, 50 mM imidazole, 1.5 M NaCl). The lysate was clarified by centrifugation at 18,000 rpm for 45 minutes at 4°C. The supernatant was loaded onto HisTrap HP column (Cytiva) for affinity purification. Protein was eluted with an imidazole gradient buffer. The elute was diluted to lower the NaCl concentration to around 0.3 M before being loaded onto HiTrap Heparin HP column (Cytiva) and eluted using a linear gradient of increasing NaCl concentration from 0.3 M to 2.0 M. The collected elute was concentrated and loaded onto a Superdex 200 Increase 10/300 column (Cytiva) pre-equilibrated with SEC buffer (20 mM Tris-HCl pH 8.0, 500 mM NaCl, 0.5 mM TCEP and 5 mM MgCl2). Peak fractions were collected and examined by SDS-PAGE. Protein samples were flash-frozen in liquid nitrogen and stored at − 80 °C.

For gRNA production, the linearized plasmid encoding gRNA sequence was used as template for in vitro transcription using HiScribe T7 High Yield RNA Synthesis Kit (New England Biolabs) following the manufacturer’s instructions. gRNA was purified by using Monarch RNA Cleanup Kit (New England Biolabs) and examined by TBE-Urea gel electrophoresis in a 15% Urea denaturing gel (Biorad).

### Cryo-EM sample preparation, data collection and processing

Prior to complex assembly, gRNA was heated at 95 °C for 2 min followed by cooling at room temperature, while the dsDNA duplex was formed by mixing equimolar targeting and non-targeting strand, heated at 95 °C for 2 min, then cooling at room temperature. Cas12f-gRNA binary complex was assembled by mixing purified cas12f protein and gRNA at a molar ratio of 1:1.2 followed by incubation at 37 °C for 30 min. dsDNA duplex (the same amount as gRNP) was then incubated with the binary complex at room temperature for 30 min to form cas12f-gRNA-DNA ternary complex. 7μM ternary complex was applied onto the Quantifoil R1.2/1.3 400 mesh copper grid that has been plasma-cleaned for 1 min by a Solarus 950 plasma cleaner (Gatan). Grids were blotted by a Vitrobot Mark IV (Thermo Fisher) for 6 s with blot force of 0 at 4 °C and 100% humidity, and plunge-frozen in liquid ethane. Grids were stored in liquid nitrogen before screening.

Grids were screened with SerialEM ^39^ on a FEI Glacios cryo-TEM. For Cas12f-MG119-28 and OsCas12f ternary complexes, images were collected on a FEI Glacios cryo-TEM equipped with a Falcon 4 detector with a pixel size of 0.933 Å, while the images of the RhCas12f ternary complex were collected on a FEI Titan Krios cryo-TEM equipped with a Gatan K3 direct electron detector with a pixel size of 0.8332 Å. The defocus range was set to -1.5 to -2.5 μm. Motion correction, contrast transfer function (CTF) estimation and particle picking were carried out in cryoSPARC live v4.0. All subsequent data processing was carried out in cryoSPARC v4.4 ^40^.

For Cas12f-MG119-28, 11,080 movies were collected, and 2,039,236 particles were extracted for downstream analysis. Particles were classified using 2D classification and classes with high resolution were picked for ab-initio reconstruction. The resulting particles and volumes were subjected to one round of heterogeneous refinement. The best class was then refined through non-uniform refinement yielding a 3.06 Å structure comprised of 507,067 particles. To sort out the heterogeneity in the dataset, these particles were subjected to a single round of 3D classification. Classes with and without a visible RuvC domain were combined and particles were re-extracted and re-refined to form the final structures (**Suppl. Fig. 6, Suppl. Fig. 7**).

For OsCas12f, 363,046 particles were picked while 223,961 particles were selected from 2D classification for further ab-initio reconstruction (3 classes). The resulting particles and volume classes were subjected to heterogeneous and non-uniform refinement which yielded a subset of 132,962 particles and a volume at 3.45 Å. The particles were re-extracted in box size of 320 pixels followed by non-uniform refinement and global/local CTF refinement which improved volume resolution to 3.27 Å that was used for modeling. Particles were subjected to 3D classification, resulting three structures (State I, State II and State III) with different R-loop formation states and RuvC.2 availability (**Suppl. Fig. 6, Suppl. Fig. 7**).

Data processing of RhCas12f ternary was performed similarly as described for the OsCas12f. 875,624 particles were selected from 2D classification for ab-initio reconstruction (3 classes). The resulting particles and volume classes were subjected to heterogeneous and non-uniform refinement which yielded a subset of 316,144 particles and a volume at 3.44 Å. The particles were re-extracted in box size of 320 pixels followed by non-uniform refinement which yielded a 3.07 Å volume that was used for modeling (**Suppl. Fig. 6, Suppl. Fig. 7**).

### Model building and refinement

The Cas12f protein structure predicted by AlphaFold2 ^41^ was fitted into the ternary complex map as a rigid body in ChimeraX ^42^. gRNA was modeled based on secondary structure prediction and gRNA architecture of AsCas12f. The model was manually refined in COOT ^43^, and automatically refined by real_space_refine in PHENIX ^44^. All structural figures were generated using ChimeraX v1.7.1. Cas12f-MG119-28 was built using Cosmic2 ModelAngelo ^45,46^. The model was subsequently manually refined using COOT and Isolde ^47^, and automatically refined by real_space_refine in PHENIX.

### GFP depletion assay

Cas12f coding sequence and DNA template of gRNA were cloned into *araBAD*-driven plasmids with resistance to chloramphenicol and carbenicillin, respectively. The GFP-encoding plasmid with streptomycin resistance was used as the target. 5 ul of *E.coli* DH5α overnight culture bearing these three plasmids were subcultured to 2.5 ml fresh LB medium. The culture was then split into two equal volumes: (1) as an uninduced control and (2) induced with 20 mM L-arabinose. All cultures were distributed into a 96-well plate (Invitrogen), then incubated at 37 °C at 300 rpm. GFP fluorescence was monitored overnight by a CLARIOstar Plus plate reader (BMG Labtech). Each measurement was repeated three times. Fold reduction was calculated by comparing fluorescence between control and induced samples.

### *In vivo* plasmid interference assay

*E.coli* cells bearing three plasmids (Cas12f-encoding, gRNA-encoding, and target plasmid) were grown up to an OD600 of approximately 0.6 at 37°C. The culture was then split into two equal volumes: (1) as an uninduced control and (2) induced with 20 mM L-arabinose. All cell cultures were incubated at 16 °C overnight, and subsequently serially diluted (six series of tenfold dilution) in fresh LB medium and spotted as 4 μl drops on LB agar plates containing all three antibiotics. LB agar plates were incubated at 37 °C overnight. Colonies were counted in the highest countable spot in the dilution series to calculate the CFU/ml. The results were compared between control and induced sample to calculate the fold reduction due to nuclease-driven plasmid cleavage. Each experiment was repeated three times as biological replicates.

### DNA cleavage kinetics

To obtain the active fraction of the enzyme an active site titration assay was performed by incubating increasing concentrations of protein relative to a fixed concentration of DNA. Briefly, pre-assembled RNP was incubated at different ratios relative to a 10nM DNA substrate in 10x Effector Buffer (100mM NaCl, 10mM Tris, 10mM MgCl2) and intubated at 37°C or 46°C for 60 minutes. For analysis, 2 uL of product was mixed with 10 uL of highly deionized (HiDi) formamide, then resolved and quantified using an Applied Biosystems DNA sequencer (ABI 3130xl) equipped with a 36 cm capillary array and nanoPOP6 polymer. The percent of cleaved DNA was evaluated by quantifying the ratio of the cleaved product relative to the uncleaved product for each concentration. All concentrations were plotted, and the active fraction was determined by fitting a line though all datapoints using a standard y-mx+b fit where b=0.

For time course assays, reactions were initiated by adding 20 nM 6-FAM labeled DNA substrate (for Os and RhCas12f and 10nM for Cas12f-MG119-28) into pre-assembled Cas12f- gRNA complex (100 nM active-site concentration of Cas12f), in 1x Effector Buffer, then quenched at various times by mixing with equal volume of 0.5 M EDTA. Reactions were performed at 46 °C for Os- and RhCas12f and at 37°C for Cas12f-MG119-28. 2 uL of product was mixed with 10 uL of highly deionized (HiDi) formamide, then resolved and quantified using an Applied Biosystems DNA sequencer (ABI 3730xl) equipped with a 36 cm capillary array and nanoPOP-7 polymer.

### Stopped-flow kinetic assay

Stopped-flow experiments were performed as previously described with minor modifications. In this experiment, position 16 in the non-target strand of the spacer portion of the DNA was labeled with the fluorophore 1,3-diaza-2-oxophenoxazine (tC°) so that upon R-loop completion, there is an increase in fluorescence as the fluorophore goes from a quenched state in the double-stranded form in target DNA to a fluorescent state when it becomes single stranded as the R-loop is formed. 100-150 nM active site Cas12f-gRNA complex was mixed with 100 nM tC°-labeled DNA substrate at 46 or 37 °C using AutoSF-120 stopped-flow instrument (KinTek Corporation, Austin, TX). Excitation was at 367 nm, and emission was monitored with a 445 nm filter with a 20 nm bandpass (Semrock).

### Global fitting of kinetic data

Stopped flow and cleavage kinetic data were globally fit to the model shown in Figure 6 with KinTek Explorer software version 11 (KinTek Corporation, Austin, TX). Rates shown in black were locked at the values listed in the fitting while other parameters were allowed to float. For parameters where only an equilibrium constant and a lower limit on a rate was obtained, the parameter with a lower limit listed in Figure 7 was locked at the lower limit in the fitting and the other rate constant for that step was allowed to float. The output for experiments to measure cleavage were modeled simply as the sum of products for each cleavage. For example, for target strand cleavage the products were defined according to our kinetic model as the sum of all species containing P2: EDP2+EDP1P2. Non-target strand cleavage products were defined as the sum of all species containing P1: EDP1+XDP1 + EDP1P2. The fluorescence signal for R-loop formation was modeled as a weighted sum of all species contributing to net fluorescence. Specifically, the fluorescence output was modeled as a*(D+ED+ EDx+ b*(EDR + EDP2 + EDP1 + EDP1P2 + XDP1)). The scaling factors, a and b, were derived as variable parameters in globally fitting the data. FitSpace confidence contour analysis was performed to define the lower and upper limits for each kinetic parameter and to establish that all parameters (**Suppl. Fig. 13**), including scaling factors, were well constrained according to this minimal model.

### Data availability

Structures of the Cas12f-MG119-28 State I, State II, OsCas12f State I, State II, State III, RhCas12f have been deposited in the EMDB with accession codes: EMD-49954, EMD-49957, EMD-49959, EMD-49956, EMD-49958, EMD-49955, respectively. Associated atomic coordinates were deposited to PDB with accession codes: 9NZO, 9NZR, 9NZT, 9NZQ, 9NZS, 9NZP, respectively. Protein and guide RNA sequences for nucleases reported here are available in Supplemental Data tables.

Structures and atomic coordinates can be found at: https://figshare.com/s/7dd7d98f6d2ef731890d.

## Acknowledgements

We thank Dr Jack Bravo for help in cryo-EM data collection and processing, thank Taylor lab members for valuable discussion, Dr. Axel Brilot and Dr. Evan Schwartz at the Sauer Lab at UT Austin for cryo-EM assistance. We also thank Dr. Emily Thompson for her thoughtful comments on the manuscript.

## Author contributions

K.G. and R.F.O performed the cryo-EM data preparation, collection, structural determination and analysis, also performed biochemical, kinetic, and in vivo assays. P.B.M.C and C.J.C performed metagenomic discovery and phylogenetic characterization. L.G.-O., L.M.A., and D.T.C. designed and performed in vitro assays. L.G.-O., M.B., and R.C.L. performed and analyzed the in vivo experiments. D.T.C. purified the proteins. L.M.A. analyzed bioinformatic data. M.M.H., N.A., and I.K. assisted with the biochemistry, kinetics and in vivo assays. T.L.D. and K.A.J performed and interpreted kinetic analysis and global fitting. K.G., R.F.O, P.B.M.C. and T.L.D. wrote the manuscript. D.W.T., D.S.A.G, C.N.B., L.M.A. and C.T.B. supervised the project, reviewed and edited the manuscript. D.W.T. and C.T.B. and provided resources and funding for the work. All authors reviewed, edited, and approved the manuscript.

## References

1. Cong, L. et al. Multiplex Genome Engineering Using CRISPR/Cas Systems. Science 339, 819–823 (2013).

2. Hwang, W. Y. et al. Efficient genome editing in zebrafish using a CRISPR-Cas system. Nat Biotechnol 31, 227–229 (2013).

3. Feng, Z. et al. Efficient genome editing in plants using a CRISPR/Cas system. Cell Res 23, 1229–1232 (2013).

4. Jinek, M. et al. A Programmable Dual-RNA–Guided DNA Endonuclease in Adaptive Bacterial Immunity. Science 337, 816–821 (2012).

5. Ran, F. A. et al. Genome engineering using the CRISPR-Cas9 system. Nat Protoc 8, 2281– 2308 (2013).

6. Koonin, E. V., Gootenberg, J. S. & Abudayyeh, O. O. Discovery of Diverse CRISPR-Cas Systems and Expansion of the Genome Engineering Toolbox. Biochemistry 62, 3465–3487 (2023).

7. Zetsche, B. et al. Cpf1 Is a Single RNA-Guided Endonuclease of a Class 2 CRISPR-Cas System. Cell 163, 759–771 (2015).

8. Gier, R. A. et al. High-performance CRISPR-Cas12a genome editing for combinatorial genetic screening. Nat Commun 11, 3455 (2020).

9. Tang, K. et al. Cas12a-knock-in mice for multiplexed genome editing, disease modelling and immune-cell engineering. *Nat*. Biomed. Eng (2025) doi:10.1038/s41551-025-01371-2.

10. Ma, E. et al. Improved genome editing by an engineered CRISPR-Cas12a. Nucleic Acids Research 50, 12689–12701 (2022).

11. Fuentes, C. M. & Schaffer, D. V. Adeno-associated virus-mediated delivery of CRISPR- Cas9 for genome editing in the central nervous system. Current Opinion in Biomedical Engineering 7, 33–41 (2018).

12. Altae-Tran, H. et al. The widespread IS200/IS605 transposon family encodes diverse programmable RNA-guided endonucleases. Science 374, 57–65 (2021).

13. Karvelis, T. et al. PAM recognition by miniature CRISPR–Cas12f nucleases triggers programmable double-stranded DNA target cleavage. Nucleic Acids Research 48, 5016– 5023 (2020).

14. Aliaga Goltsman, D. S., et al. Compact Cas9d and HEARO enzymes for genome editing discovered from uncultivated microbes. Nat Commun 13, 7602 (2022).

15. Harrington, L. B. et al. Programmed DNA destruction by miniature CRISPR-Cas14 enzymes. Science 362, 839–842 (2018).

16. Bigelyte, G. et al. Miniature type V-F CRISPR-Cas nucleases enable targeted DNA modification in cells. Nat Commun 12, 6191 (2021).

17. Kong, X. et al. Engineered CRISPR-OsCas12f1 and RhCas12f1 with robust activities and expanded target range for genome editing. Nat Commun 14, 2046 (2023).

18. Xiao, R., Li, Z., Wang, S., Han, R. & Chang, L. Structural basis for substrate recognition and cleavage by the dimerization-dependent CRISPR–Cas12f nuclease. Nucleic Acids Research 49, 4120–4128 (2021).

19. Takeda, S. N. et al. Structure of the miniature type V-F CRISPR-Cas effector enzyme. Molecular Cell 81, 558–570.e3 (2021).

20. Wu, T. et al. An engineered hypercompact CRISPR-Cas12f system with boosted gene-editing activity. Nat Chem Biol 19, 1384–1393 (2023).

21. Wu, Z. et al. Structure and engineering of miniature Acidibacillus sulfuroxidans Cas12f1. Nat Catal 6, 695–709 (2023).

22. Su, M. et al. Molecular basis and engineering of miniature Cas12f with C-rich PAM specificity. Nat Chem Biol (2023) doi:10.1038/s41589-023-01420-4.

23. Hino, T. et al. An AsCas12f-based compact genome-editing tool derived by deep mutational scanning and structural analysis. Cell S0092867423009637 (2023) doi:10.1016/j.cell.2023.08.031.

24. Wu, Z. et al. Programmed genome editing by a miniature CRISPR-Cas12f nuclease. Nat Chem Biol 17, 1132–1138 (2021).

25. Zhang, X. et al. Engineered circular guide RNAs enhance miniature CRISPR/Cas12f- based gene activation and adenine base editing. Nat Commun 16, 3016 (2025).

26. Stella, S. et al. Conformational Activation Promotes CRISPR-Cas12a Catalysis and Resetting of the Endonuclease Activity. Cell 175, 1856–1871.e21 (2018).

27. Zhang, H., Li, Z., Xiao, R. & Chang, L. Mechanisms for target recognition and cleavage by the Cas12i RNA-guided endonuclease. Nat Struct Mol Biol 27, 1069–1076 (2020).

28. Kurihara, N. et al. Structure of the type V-C CRISPR-Cas effector enzyme. Molecular Cell 82, 1865–1877.e4 (2022).

29. Gong, S., Yu, H. H., Johnson, K. A. & Taylor, D. W. DNA Unwinding Is the Primary Determinant of CRISPR-Cas9 Activity. Cell Reports 22, 359–371 (2018).

30. 30. Liu, M.-S., Gong, S., Yu, H.-H., Taylor, D. W. & Johnson, K. A. Chapter Thirteen - Kinetic characterization of Cas9 enzymes. in Methods in Enzymology (ed. Bailey, S.) vol. 616 289–311 (Academic Press, 2019).

31. 31. Johnson, K. A. Chapter 23 Fitting Enzyme Kinetic Data with KinTek Global Kinetic Explorer. in Methods in Enzymology vol. 467 601–626 (Elsevier, 2009).

32. Johnson, K. A., Simpson, Z. B. & Blom, T. FitSpace Explorer: An algorithm to evaluate multidimensional parameter space in fitting kinetic data. Analytical Biochemistry 387, 30–41 (2009).

33. Ocampo, R. F. et al. DNA targeting by compact Cas9d and its resurrected ancestor. Nat Commun 16, 457 (2025).

34. Liu, M.-S. et al. Engineered CRISPR/Cas9 enzymes improve discrimination by slowing DNA cleavage to allow release of off-target DNA. Nat Commun 11, 3576 (2020).

35. Bravo, J. P. K. et al. Structural basis for mismatch surveillance by CRISPR–Cas9. Nature 603, 343–347 (2022).

36. Pacesa, M. et al. R-loop formation and conformational activation mechanisms of Cas9. Nature 609, 191–196 (2022).

37. Li, D., Liu, C.-M., Luo, R., Sadakane, K. & Lam, T.-W. MEGAHIT: an ultra-fast single- node solution for large and complex metagenomics assembly via succinct *de Bruijn* graph. Bioinformatics 31, 1674–1676 (2015).

38. Hyatt, D. et al. Prodigal: prokaryotic gene recognition and translation initiation site identification. BMC Bioinformatics 11, 119 (2010).

39. Mastronarde, D. N. Automated electron microscope tomography using robust prediction of specimen movements. Journal of Structural Biology 152, 36–51 (2005).

40. Punjani, A., Rubinstein, J. L., Fleet, D. J. & Brubaker, M. A. cryoSPARC: algorithms for rapid unsupervised cryo-EM structure determination. Nat Methods 14, 290–296 (2017).

41. Jumper, J. et al. Highly accurate protein structure prediction with AlphaFold. Nature 596, 583–589 (2021).

42. Pettersen, E. F. et al. UCSF CHIMERAX : Structure visualization for researchers, educators, and developers. Protein Science 30, 70–82 (2021).

43. Emsley, P., Lohkamp, B., Scott, W. G. & Cowtan, K. Features and development of *Coot*. Acta Crystallogr D Biol Crystallogr 66, 486–501 (2010).

44. Liebschner, D. et al. Macromolecular structure determination using X-rays, neutrons and electrons: recent developments in *Phenix*. Acta Crystallogr D Struct Biol 75, 861–877 (2019).

45. Cianfrocco, M. A., Wong-Barnum, M., Youn, C., Wagner, R. & Leschziner, A. COSMIC2: A Science Gateway for Cryo-Electron Microscopy Structure Determination. in Practice and Experience in Advanced Research Computing 2017: Sustainability, Success and Impact 1–5 (ACM, New Orleans LA USA, 2017). doi:10.1145/3093338.3093390.

46. Jamali, K. et al. Automated model building and protein identification in cryo-EM maps. Nature 628, 450–457 (2024).

47. Croll, T. I. *ISOLDE* : a physically realistic environment for model building into low- resolution electron-density maps. Acta Crystallogr D Struct Biol 74, 519–530 (2018).

