## Supplementary Information for "Comparative characterization of Cas12f orthologs reveals mechanistic features underlying enhanced genome editing efficiency"

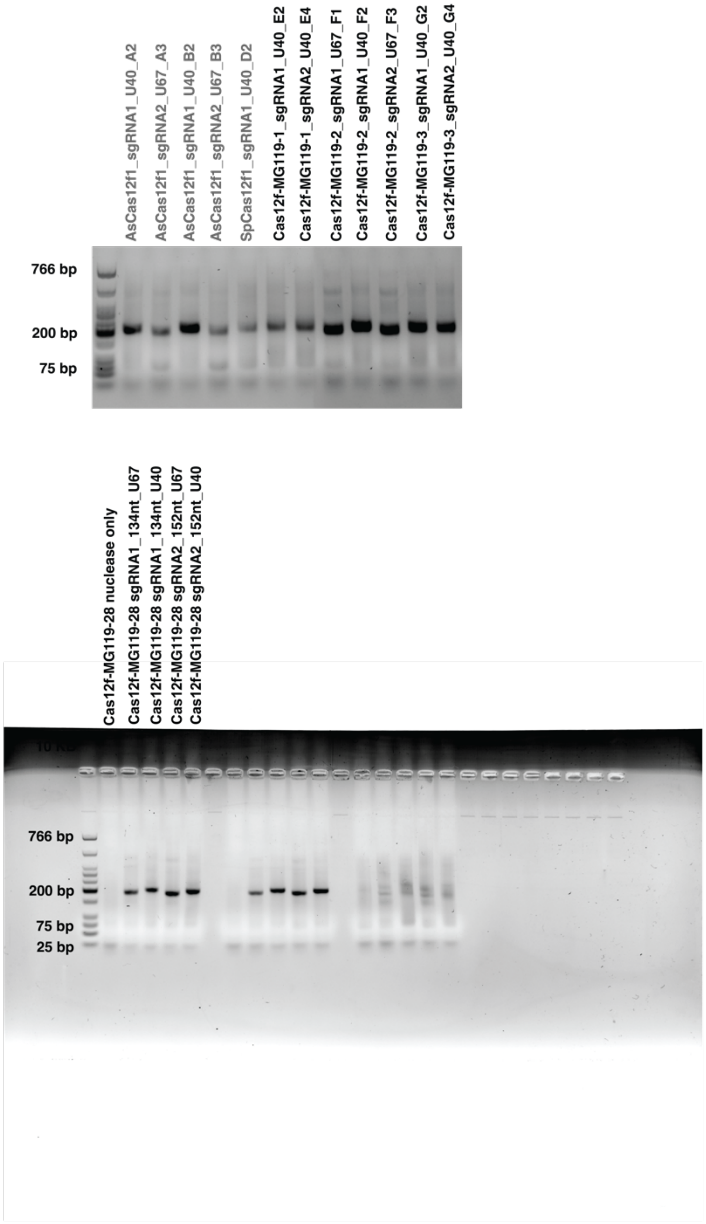


**Supplementary Figure 1. Validation of the sgRNA designs.**

Cleavage activity was detected on 2% agarose gels (NEB Low Molecular Weight DNA Ladder). Publicly available systems (AsCas12f1 and SpCas12f1) were added for reference. Cleavage activity for each system was tested with a couple of predicted sgRNAs.

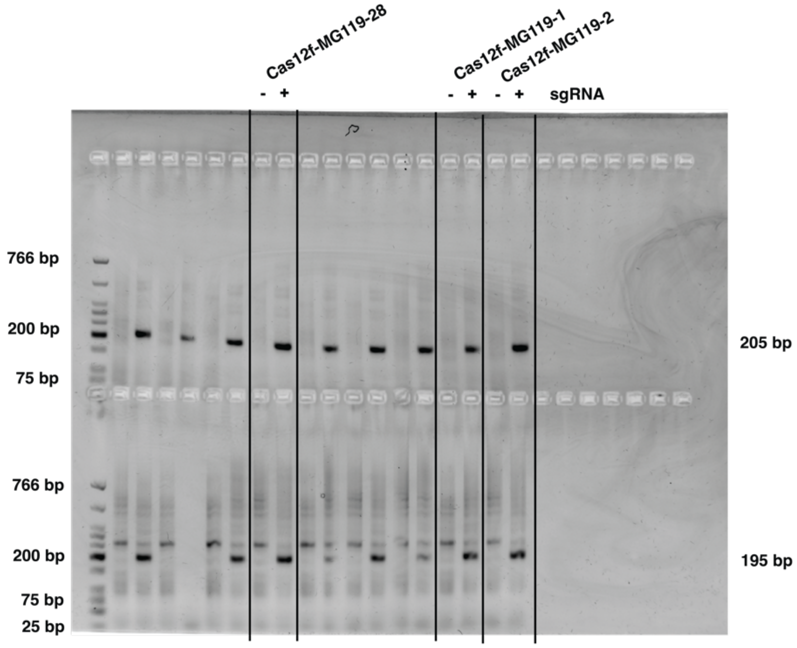


**Supplementary Figure 2. Double strand cleavage products.**

Agarose gel (2%) of amplified cleavage products to capture the target strand cut site (top) and the non-target strand cut site (bottom). Left lane (-): nuclease only. Right lane (+): nuclease plus sgRNA (NEB Low Molecular Weight DNA Ladder).

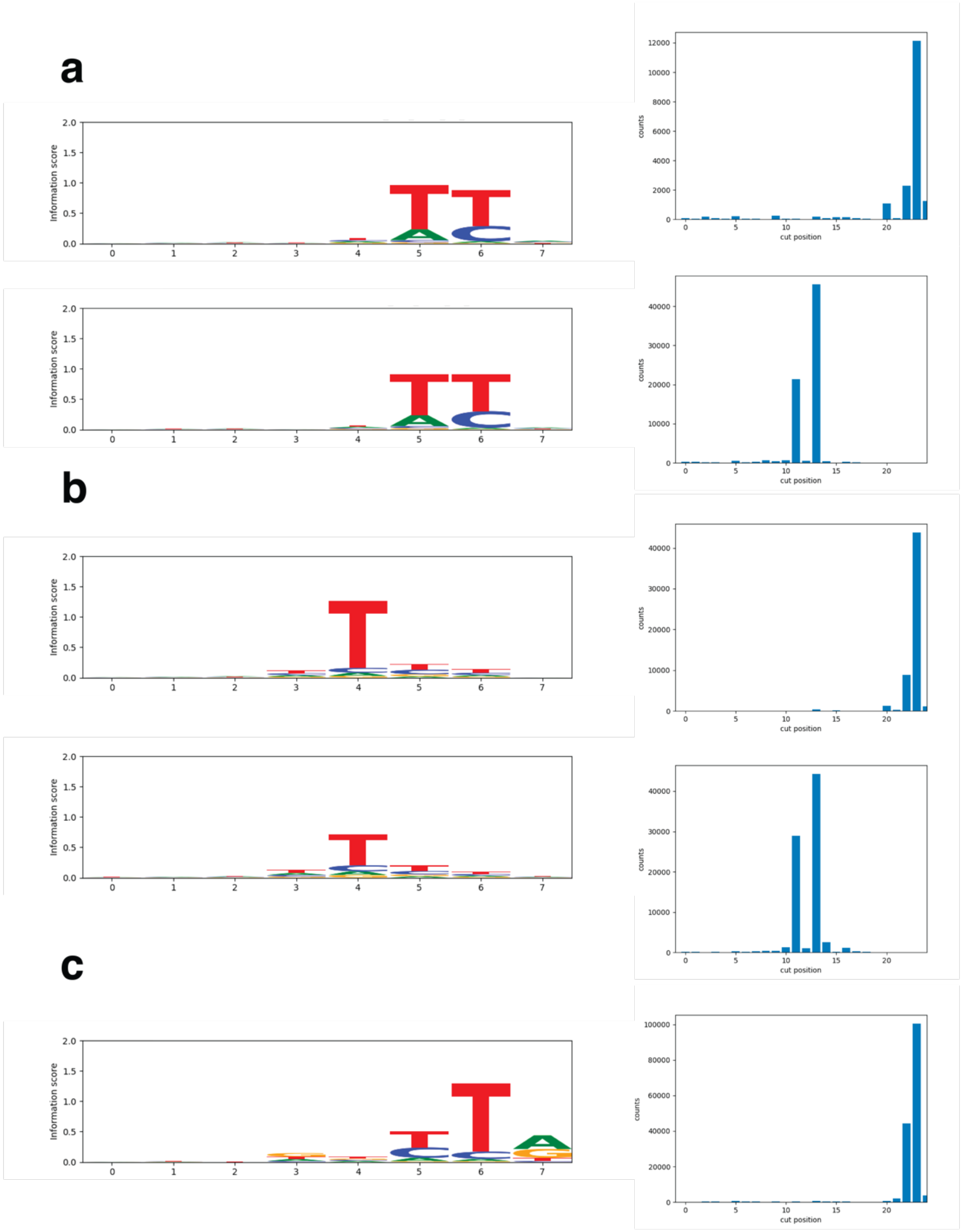


**Supplementary Figure 3. SeqLogos and cut site histograms.**

**a.** Cas12f-MG119-1. **b.** Cas12f-MG119-2. **c.** Cas12f-MG119-3. The PAM sequences were confirmed by NGS and represented as SeqLogos (Top: target strand; Bottom: non-target strand). Histograms show the number of NGS reads (counts) mapped to the cleavage site on each strand. Cas12f-MG119-3 cut site on the NTS was less than a 1000 reads (not shown).

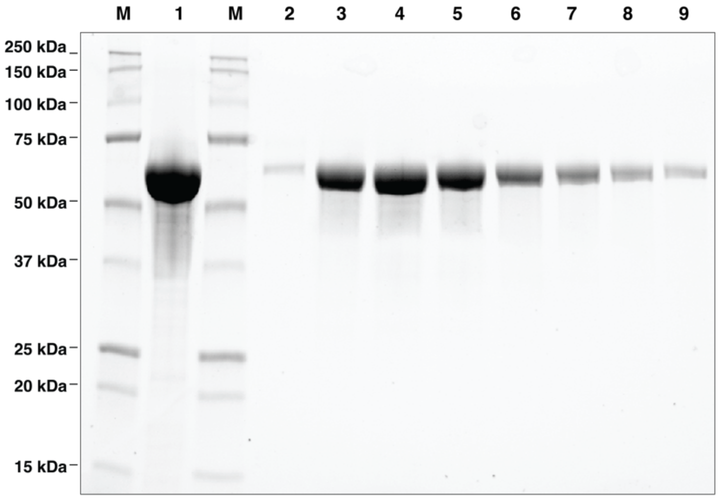


**Supplementary Figure 4. SDS-PAGE gel of Cas12f-MG119-28 protein purification steps.**

Sample before (1) and after fraction samples (2-9) SEC. M, Marker (Bio-Rad Catalog # 1610363).

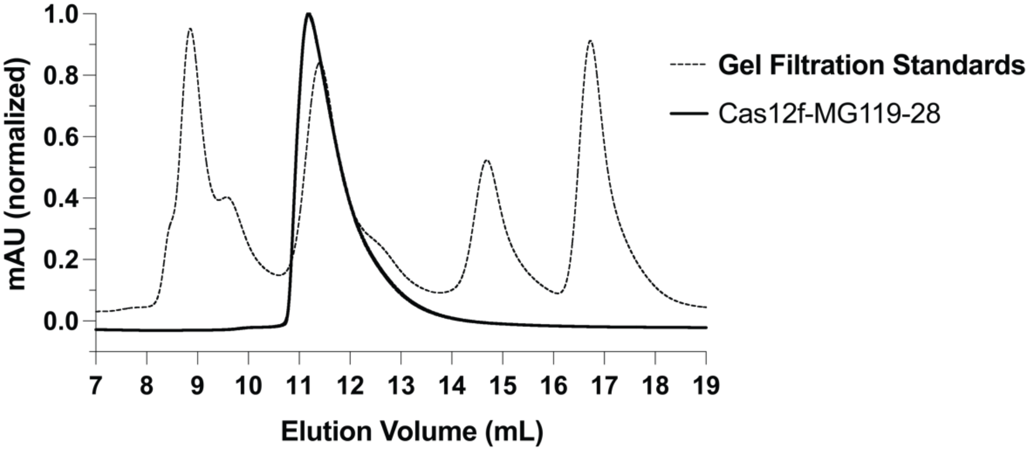


**Supplementary Figure 5. Cas12f-MG119-28 is an obligate dimer.**

SEC elution trace of Cas12f-MG119-28 (dark line) overlaid with the elution trace of BioRad Gel Filtration Standards (dashed line, Bio-Rad). The Cas12f-MG119-28 peak elutes around 11.5 mL post-injection, similar to the second peak from the gel filtration standards (bovine γ-globulin, approx. 158,000 Da).

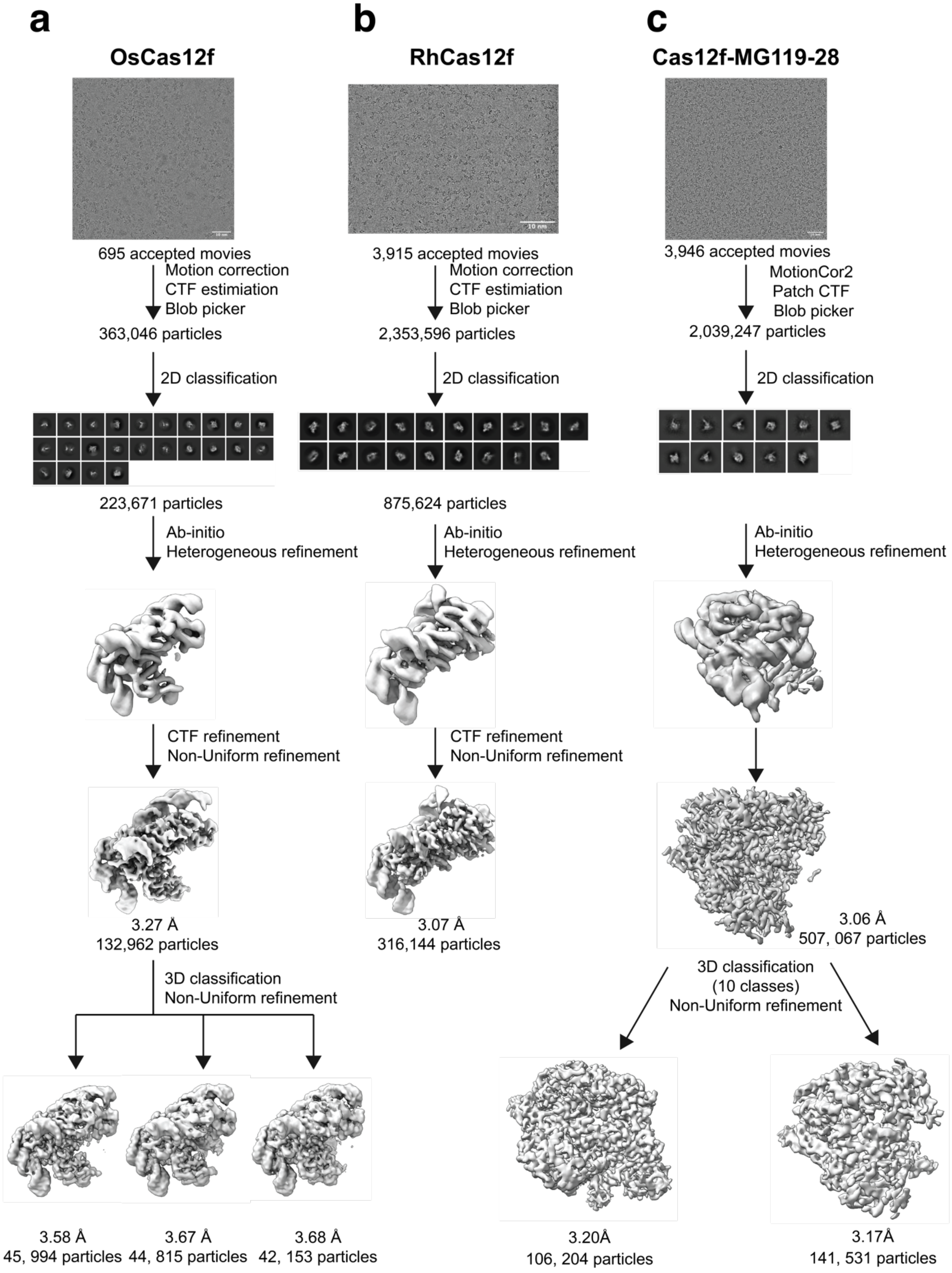

a
OsCas12f
695 accepted movies
Motion correction
CTF estiimation
Blob picker
363,046 particles
2D classification
223,671 particles
Ab-initio
Heterogeneous refinement
CTF refinement
Non-Uniform refinement
3.27 Å
132,962 particles
3D classification
Non-Uniform refinement
3.58 Å
45, 994 particles
3.67 Å
44, 815 particles
3.68 Å
42, 153 particles
b
RhCas12f
3,915 accepted movies
Motion correction
CTF estiimation
Blob picker
2,353,596 particles
2D classification
875,624 particles
Ab-initio
Heterogeneous refinement
CTF refinement
Non-Uniform refinement
3.07 Å
316,144 particles
c
Cas12f-MG119-28
3,946 accepted movies
MotionCor2
Patch CTF
Blob picker
2,039,247 particles
2D classification
Ab-initio
Heterogeneous refinement
3.06 Å
507, 067 particles
3D classification
(10 classes)
Non-Uniform refinement
3.20Å
106, 204 particles
3.17Å
141, 531 particles

**Supplementary Figure 6. Cryo-EM data processing workflow for Cas12f ternary complexes.**

**a.** OsCas12f. **b.** RhCas12f. **c.** Cas12f-MG119-28.

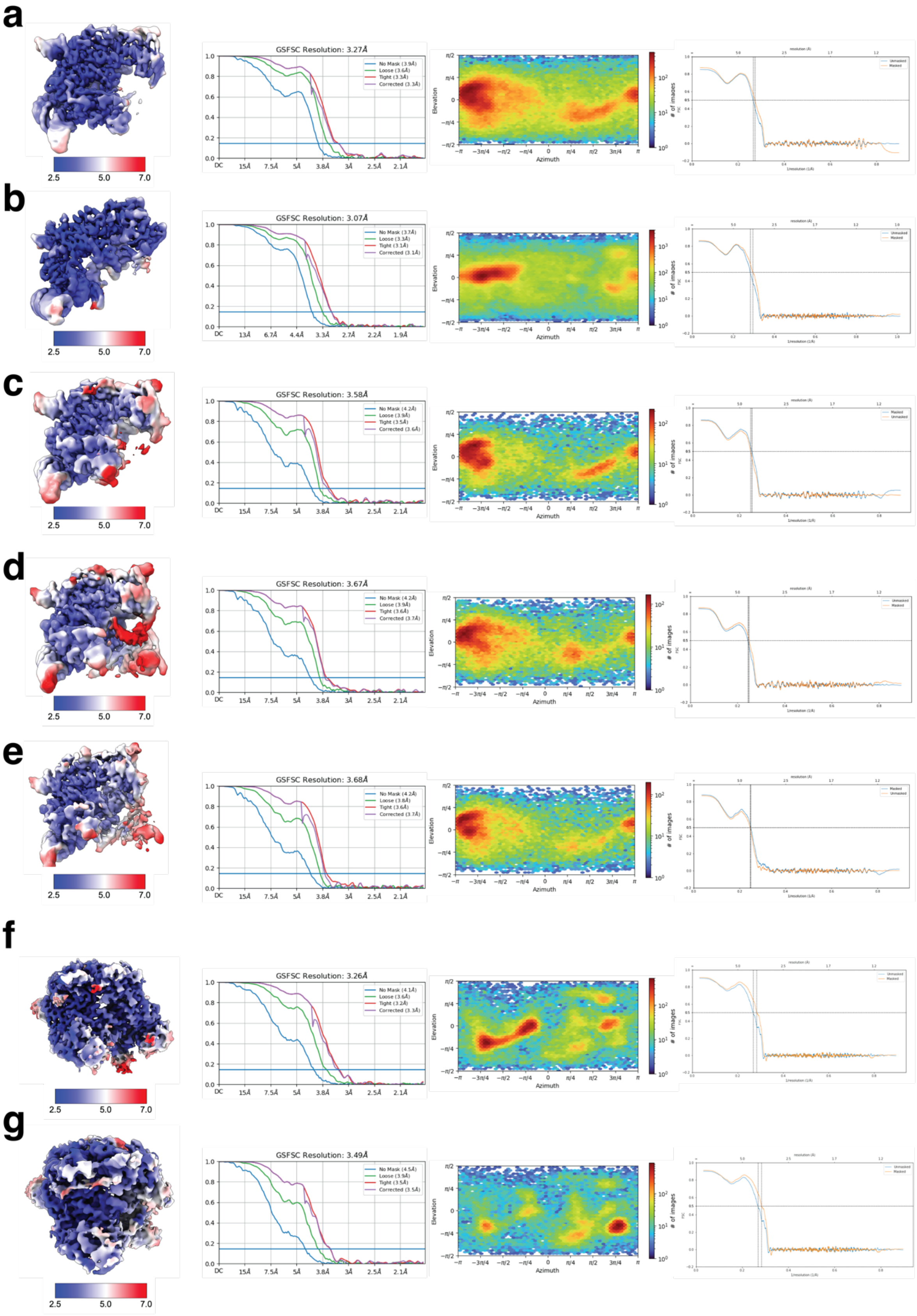

a
2.5
5.0
7.0
GSFSC Resolution: 3.27Å
No Mask (3.9Å)
Loose (3.6Å)
Tight (3.3Å)
Corrected (3.3Å)
Elevation
Azimuth
### of images
Unmasked
Masked
1/resolution (1/Å)
b
GSFSC Resolution: 3.07Å
No Mask (3.7Å)
Loose (3.3Å)
Tight (3.1Å)
Corrected (3.1Å)
c
GSFSC Resolution: 3.58Å
No Mask (4.2Å)
Loose (3.9Å)
Tight (3.5Å)
Corrected (3.6Å)
d
GSFSC Resolution: 3.67Å
No Mask (4.2Å)
Loose (3.9Å)
Tight (3.6Å)
Corrected (3.7Å)
e
GSFSC Resolution: 3.68Å
No Mask (4.2Å)
Loose (3.8Å)
Tight (3.6Å)
Corrected (3.7Å)
f
GSFSC Resolution: 3.26Å
No Mask (4.1Å)
Loose (3.6Å)
Tight (3.2Å)
Corrected (3.3Å)
g
GSFSC Resolution: 3.49Å
No Mask (4.5Å)
Loose (3.9Å)
Tight (3.5Å)
Corrected (3.5Å)

**Supplementary Figure 7. Cryo-EM data analysis of Cas12f complexes.**

For each structure (OsCas12f (**a**), RhCas12f (**b**), OsCas12f intermediate states I-III (**c-e**), Cas12f-MG119-28 state I-II (**f-g**). Left to right panels show local resolution maps; gold-standard Fourier Shell Correlation (FSC) curves with resolution reported at FSC = 0.143; Euler angle distribution plots of particle orientations; map-to-model FSC curves.

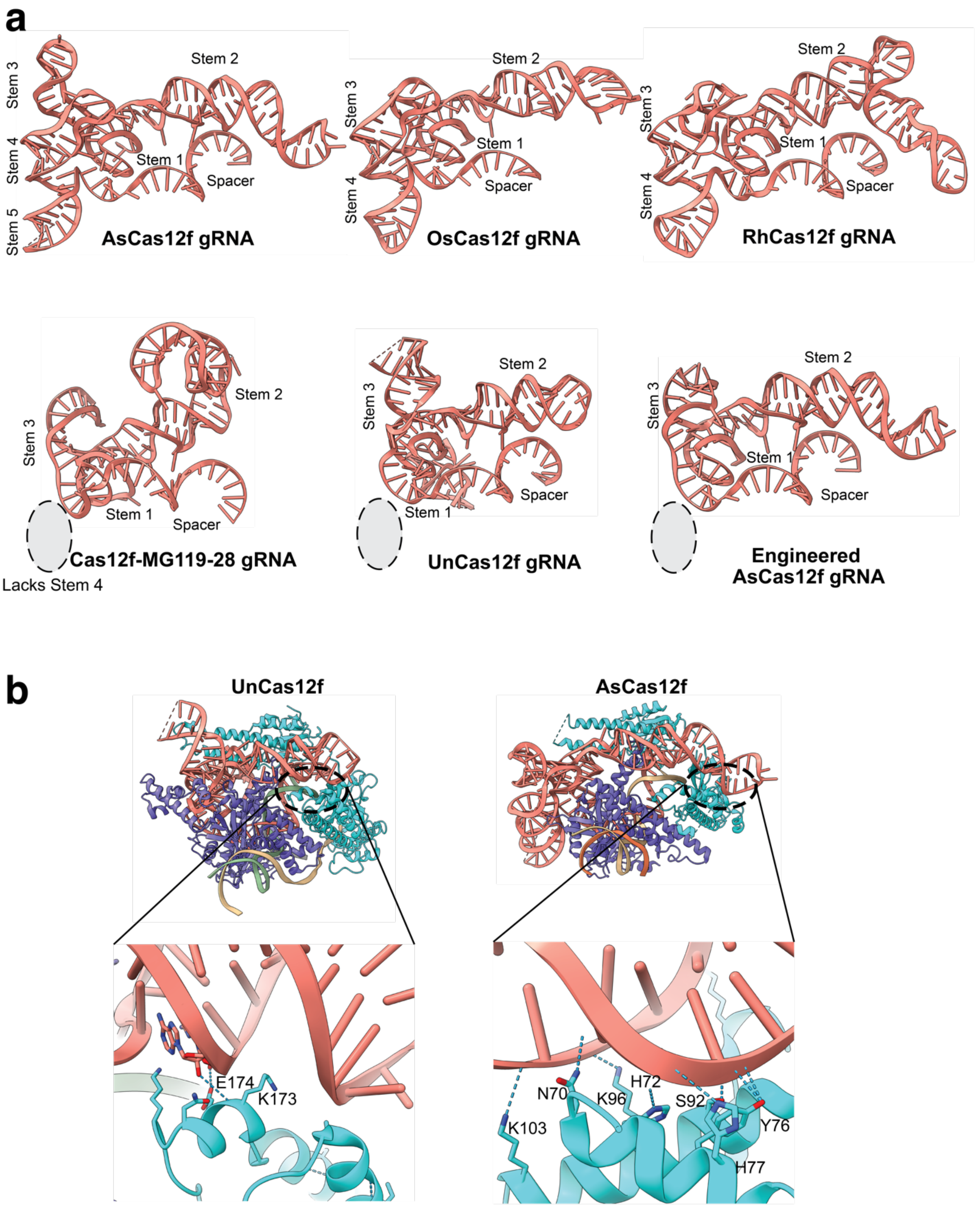

a
Stem 2
Stem 3
Stem 4
Stem 5
Stem 1
Spacer
AsCas12f gRNA
OsCas12f gRNA
RhCas12f gRNA
Cas12f-MG119-28 gRNA
Lacks Stem 4
UnCas12f gRNA
Engineered AsCas12f gRNA
b
UnCas12f
AsCas12f
E174
K173
N70
K96
H72
S92
Y76
K103
H77

**Supplementary Figure 8. Comparative analysis of gRNA architectures across Cas12f orthologs.**

**a.** gRNA scaffolds of AsCas12f (PDB: 8J12), OsCas12f, RhCas12f, Cas12f-MG119-28, UnCas12f (PDB: 7L49), and engineered AsCas12f (PDB: 8J1J). Stems, spacers, and unresolved regions (dashed lines) are labeled. **b.** Structures of UnCas12f and AsCas12f ternary complexes. Insets highlight hydrogen-bond interactions (blue dashed lines) between the gRNA Stem 2 terminus and Molecule 2.

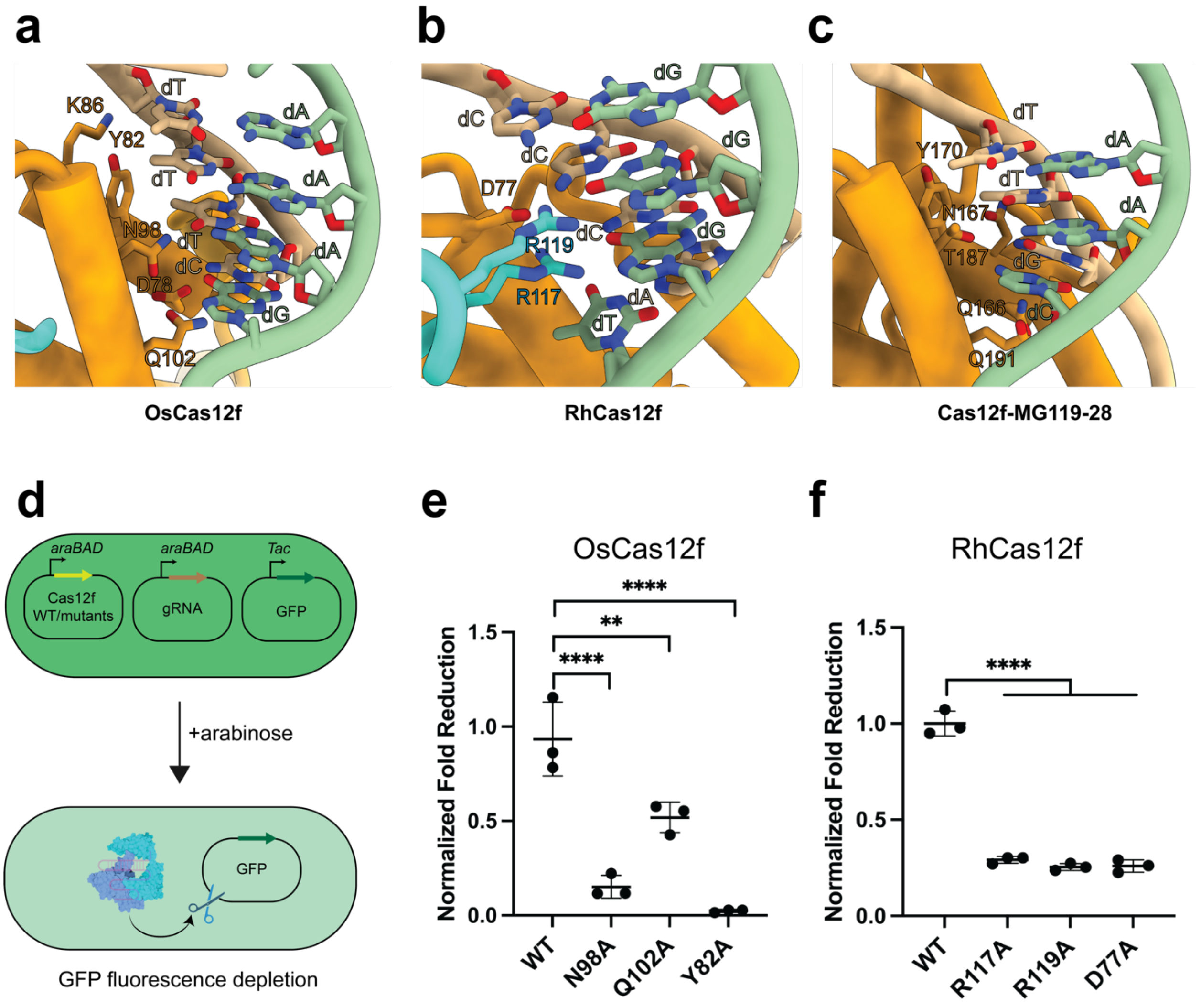


**Supplementary Figure 9. PAM recognition by Os-, RhCas12f and Cas12f-MG119-28.** Interactions between the PAM duplex and OsCas12f (**a**), RhCas12f (**b**), and Cas12f-MG119-28 (**c**). **d.** Schematic of the GFP depletion assay. e-f. In vivo activity of wild-type (WT) and PAM-recognition mutants of OsCas12f (**e**) and RhCas12f (**f**). Data represent mean ± SD (n = 3 biological replicates). Significance determined by one-way ANOVA: ***p* ≤ 0.01, *****p* ≤ 0.0001.

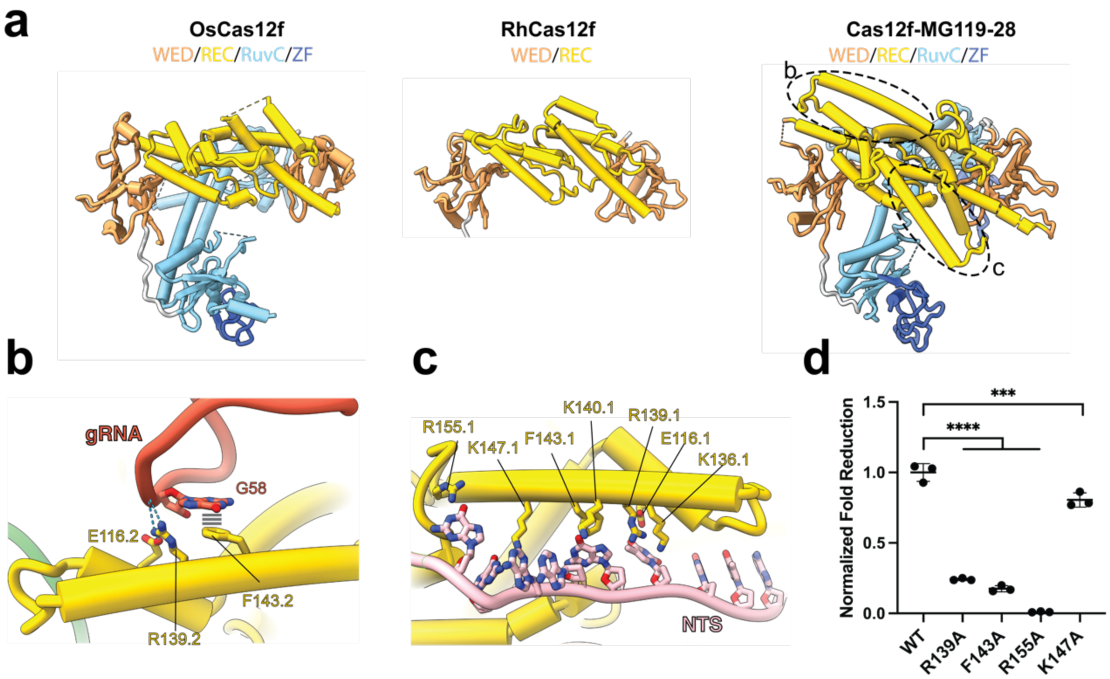


**Supplementary Figure 10. Extended helices in Cas12f-MG119-28 REC domain stabilize gRNA and non-target strand DNA.**

**a.** Protein structure comparison of Os- RhCas12f, and Cas12f-MG119-28. **b.** Interaction between the extended helix in Cas12f-MG119-28 Molecule 2 and the gRNA Stem 2 terminus. **c.** Interaction between the extended helix in Cas12f-MG-119-28 Molecule 1 and the NTS DNA. **d.** In vivo activity of WT and helix mutants of Cas12f-MG119-28 in GFP depletion assay. Data represent mean ± SD (n = 3 biological replicates). Significance determined by one-way ANOVA: ***$p \leq 0.001$, ****$p \leq 0.0001$.

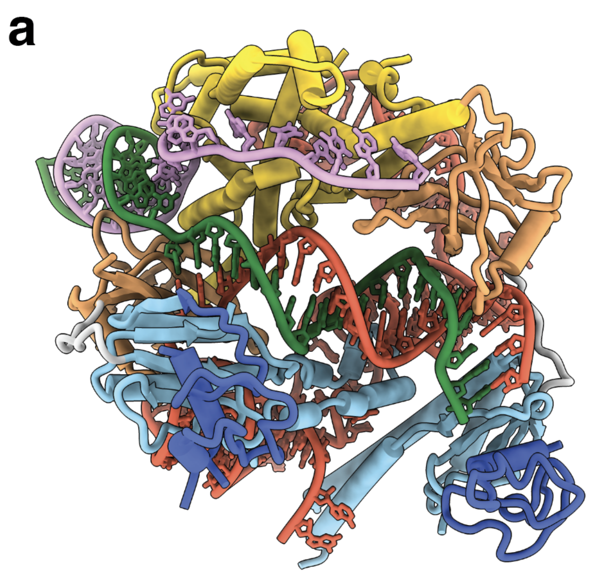

**Supplementary Figure 11. Comparison of Cas12f-MG119-28 Structures.**

**a.** Cas12f-MG119-28 structure in State II **b.** Cas12f-MG119-28 structure in State I. State I does not contain a resolved RuvC.2 domain. Instead, an additional unstructured region of the gRNA is resolved.

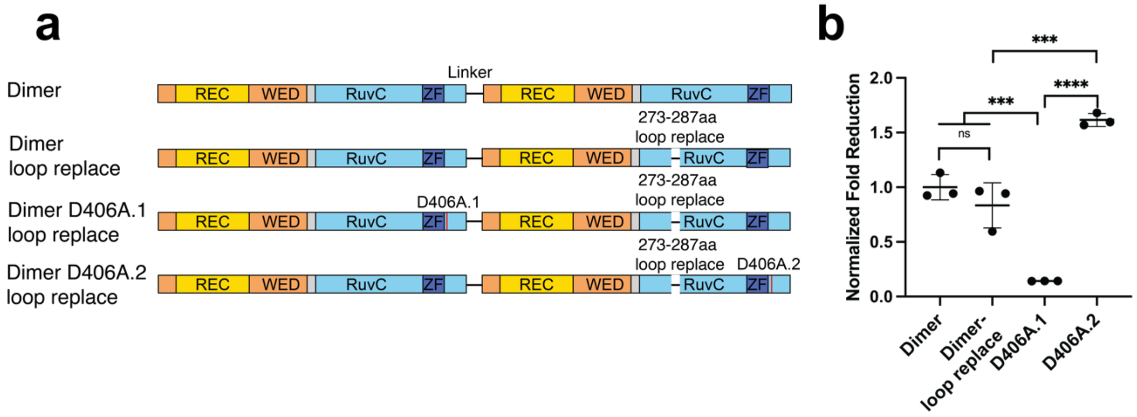


**Supplementary Figure 12. Molecule 1 mediates catalytic activity in OsCas12f.**

**a.** Design of OsCas12f covalent dimer mutants. **b.** In vivo activity of dimer WT and mutants of OsCas12f in GFP depletion assay. Data represent mean ± SD (n = 3 biological replicates). Significance determined by one-way ANOVA: ns not significant, ****p* ≤ 0.001, *****p* ≤ 0.0001.

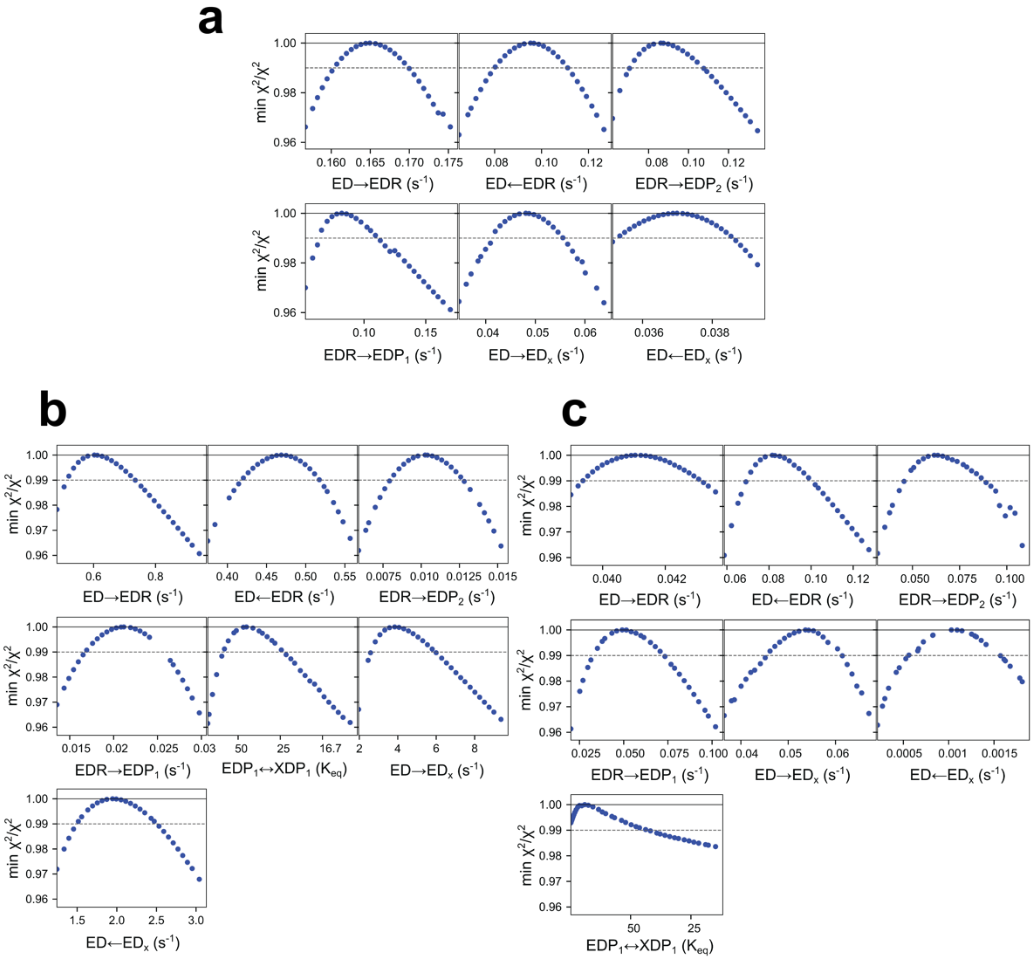


**Supplementary Figure 13. Confidence contours from global data fitting.**

**a.** Cas12f-MG119-28; **b.** OsCas12f; and **c.** RhCas12f.

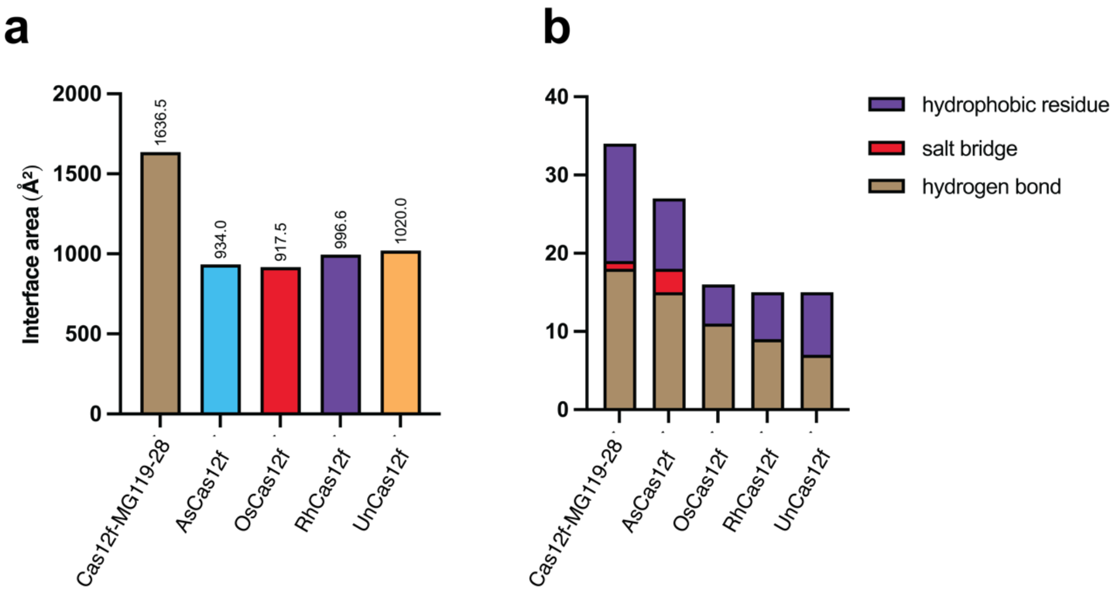


**Supplementary Figure 14. Dimer interface analysis of Cas12f nucleases.**

**a.** Interface surface area mediated by the REC domain, calculated using PDBePISA. **b.** Quantification of dimer interface interactions (hydrogen bonds, salt bridges, and hydrophobic residues). Hydrogen bonds and salt bridges were computed with PDBePISA; hydrophobic residues were manually annotated.

#### Supplementary Table 1 Cryo-EM data collection and model validation statistics

| | Cas12f-MG119-28 State I | Cas12f-MG119-28 State II | OsCas12f State I | OsCas12f State II | OsCas12f State III | RhCas12f |
|---|---|---|---|---|---|---|
| **Data collection and processing** | | | | | | |
| Magnification | 150K | 150K | 150K | 150K | 150K | 105K |
| Voltage (kV) | 200 | 200 | 200 | 200 | 200 | 300 |
| Electron exposure (e–/Å$^2$) | 50 | 50 | 50 | 50 | 50 | 80 |
| Defocus range (µm) | -1.5 to -2.5 | -1.5 to -2.5 | -1.5 to -2.5 | -1.5 to -2.5 | -1.5 to -2.5 | -1.5 to -2.5 |
| Pixel size (Å) | 0.94 | 0.94 | 0.94 | 0.94 | 0.94 | 0.83 |
| Initial particle (no.) | 2,039,247 | 2,039,247 | 223, 671 | 223, 671 | 223, 671 | 875, 624 |
| Final particle (no.) | 141,537 | 106,204 | 132, 962 | 44, 815 | 42, 153 | 316, 144 |
| Map resolution (Å) | 3.17 | 3.20 | 3.27 | 3.67 | 3.68 | 3.07 |
| FSC threshold | 0.143 | 0.143 | 0.143 | 0.143 | 0.143 | 0.143 |
| **Refinement** | | | | | | |
| Initial model used | ModelAngelo | ModelAngelo | AlphaFold | 9NZT | 9NZT | AlphaFold |
| Model resolution (Å) | 3.19 | 3.25 | 3.2 | 3.7 | 3.6 | 3.1 |
| FSC threshold | 0.143 | 0.143 | 0.143 | 0.143 | 0.143 | 0.143 |
| Map sharpening B factor (Å$^2$) | -109.1 | -117.0 | -127.7 | -107.4 | -104.3 | -135.7 |
| Model composition | | | | | | |
| Non-hydrogen atoms | 9025 | 11066 | 7928 | 9260 | 9448 | 7397 |
| Protein residues | 694 | 889 | 598 | 750 | 728 | 389 |
| Nucleotides | 169 | 185 | 146 | 152 | 170 | 199 |
| Ligands | N/A | N/A | Zn: 1 | Zn: 1 | Zn: 1 | NA |
| Mean B factors (Å$^2$) | | | | | | |
| Protein | 93.36 | 83.04 | 70.81 | 39.83 | 38.94 | 59.33 |
| Nucleotides | 164.30 | 117.10 | 94.92 | 64.73 | 67.14 | 151.79 |
| Ligands | N/A | N/A | 159.75 | 93.81 | 101.67 | NA |
| R.m.s deviations | | | | | | |
| Bond lengths (Å) | 0.005 | 0.004 | 0.005 | 0.006 | 0.007 | 0.006 |
| Bond angles (°) | 0.883 | 0.843 | 1.035 | 1.216 | 1.250 | 1.116 |
| **Validation** | | | | | | |
| MolProbity score | 1.54 | 1.32 | 1.34 | 1.53 | 1.50 | 1.35 |
| Clashscore | 5.46 | 5.86 | 4.19 | 5.79 | 7.06 | 6.39 |
| Poor rotamers (%) | 0.61 | 0.53 | 0.77 | 1.09 | 0.48 | 0.28 |
| Ramachandran plot | | | | | | |
| Favored (%) | 97.38 | 98.40 | 97.29 | 96.88 | 97.47 | 98.70 |
| Allowed (%) | 2.62 | 1.60 | 2.71 | 3.12 | 2.53 | 1.30 |
| Disallowed (%) | 0 | 0 | 0 | 0 | 0 | 0 |
| **PDB** | **9NZO** | **9NZR** | **9NZT** | **9NZQ** | **9NZS** | **9NZP** |
| **EMDB** | **EMD-49954** | **EMD-49957** | **EMD-49959** | **EMD-49956** | **EMD-49958** | **EMD-49955** |
